# Global drivers of eukaryotic plankton biogeography in the sunlit ocean

**DOI:** 10.1101/2020.09.08.287524

**Authors:** Sommeria-Klein Guilhem, Watteaux Romain, Iudicone Daniele, Bowler Chris, Morlon Hélène

## Abstract

Eukaryotic plankton are a core component of marine ecosystems with exceptional taxonomic and ecological diversity. Yet how their ecology interacts with the environment to drive global distribution patterns is poorly understood. Here, we use *Tara* Oceans metabarcoding data covering all the major ocean basins combined with a probabilistic model of taxon co-occurrence to compare the biogeography of 70 major groups of eukaryotic plankton. We uncover two main axes of biogeographic variation. First, more diverse groups display stronger biogeographic structure. Second, large-bodied consumers are structured by oceanic basins, mostly via the main currents, while small-bodied phototrophs are structured by latitude, with a comparatively stronger influence of environmental conditions. Our study highlights striking differences in biogeographies across plankton groups and investigates their determinants at the global scale.

**One-sentence summary:** Eukaryotic plankton biogeography and its determinants at global scale reflect differences in ecology and body size.

## Main text

Marine plankton communities play key ecological roles at the base of oceanic food chains, and in driving global biogeochemical fluxes (*1*, *2*). Understanding their spatial patterns of distribution is a long-standing challenge in marine ecology that has lately become a key part of the effort to model the response of oceans to environmental changes (*3*–*6*). Part of the difficulty lies in the constant recirculation of plankton communities by ocean currents, along which many physical, chemical and biological processes - the so-called *seascape* (*7*) - modify community composition (*8*). Recent planetary-scale ocean sampling expeditions have revealed that eukaryotic plankton are taxonomically and ecologically extremely diverse, possibly even more so than prokaryotic plankton (*9*). Eukaryotic plankton range from pico-sized (0.2-2 μm) to meso-sized (0.2-20 mm) organisms and larger, thus covering an exceptional range of sizes. Eukaryotic plankton also cover a wide range of ecological roles, from phototrophs (e.g., Bacillariophyta, Haptophyta, Mamiellophyceae) to parasites (e.g., Marine Alveolates or MALVs), and from heterotrophic protists (e.g., Diplonemida, Ciliophora, Acantharea) to metazoans (e.g., Arthropoda and Chordata, respectively represented principally by Copepods and Tunicates). Understanding how these body size and ecological differences modulate the influence of oceanic currents and local environmental conditions on geographic distributions is needed if one wants to predict how eukaryotic communities, and therefore the trophic interactions and global biogeochemical cycles they participate in, will change with changing environmental conditions.

Previous studies suggested that all eukaryotes up to a size of approximately 1 mm are globally dispersed and primarily constrained by abiotic conditions (*10*). While this view has been revised, the influence of body size on biogeography is manifest (*11*, *12*). In particular, a parallel study by Richter et al. (*12*), which quantified changes in plankton metagenomic composition and highlighted the underlying dynamics using transport time along main currents, found that the turnover is slower, rather than faster, with increasing body size. This suggests that, rather than influencing biogeography through its effect on abundance and ultimately dispersal capacity (i.e., larger organisms are more dispersal-limited; *10*, *11*), body size influences biogeography through its relationship with ecology and ultimately the sensitivity of communities to environmental conditions as they drift along currents. Under this scenario, the distribution of large long-lived generalist predators such as Copepods (Arthropoda) is expected to be stretched through large-scale transport by main currents (*8*, *12*–*14*), and yet to be patchy as a result of small-scale turbulent stirring (*15*). These contrasted views illustrate that little is known on how the interplay between body size, ecology, currents and the local environment shapes biogeography (*16*).

Here, we study plankton biogeography across all major eukaryotic groups in the sunlit ocean using 18S rDNA metabarcoding data from the *Tara* Oceans global survey (including recently released data from the Arctic Ocean; *17*). We also used transport times from Richter et al. (*12*), and the same environmental data. The data encompass 250,057 eukaryotic Operational Taxonomic Units (OTUs) sampled globally at the surface and at the Deep Chlorophyl Maximum (DCM) across 129 stations. We use a probabilistic model that allows identification of a number of ‘assemblages’, each of which represents a set of OTUs that tend to co-occur across samples (*18*, *19*; cf. Mat. & Meth.). Each local planktonic community can then be seen as a sample drawn in various proportions from the assemblages. Across the *Tara* Oceans samples and considering all eukaryotic OTUs together, we identified 16 geographically structured assemblages, each composed of OTUs covering the full taxonomic range of eukaryotic plankton (Fig. 1, S1; Appendix). Local planktonic communities often cannot be assigned to a single assemblage, as would be typical for terrestrial macro-organisms on a fixed landscape (*20*, *21*), but are instead mixtures of assemblages (Fig. 1A). This is consistent with previous findings suggesting that neighbouring plankton communities are continuously mixed and dispersed by currents (*8*, *12*). Nevertheless, three assemblages are particularly represented and most communities are dominated by one of them (Fig. 1A). The most prevalent assemblage represents a set of OTUs (about one fifth of the total) that are globally ubiquitous except in the Arctic Ocean (assemblage 1, in dark red). This assemblage typically accounts for about half the number of OTUs in non-Arctic communities, and is particularly rich in parasitic groups such as MALV (Fig. 1B). The two others dominate, respectively, in the Arctic Ocean (assemblage 13, in cyan) and in the Southern Ocean (assemblage 15, in marine blue), and are particularly rich in diatoms (Fig. 1B). Based on similarity in their OTU composition, the assemblages cluster into three main categories corresponding to low, intermediate and high latitudes (Fig. 1B). The transition between communities composed of high-latitude and lower-latitude assemblages is fairly abrupt, and occurs around 45° in the North Atlantic and −47° in the South Atlantic, namely at the latitude of the subtropical front, where the transition between cold and warm waters takes place (Fig. 1A&B; *22*).

**Fig. 1.**
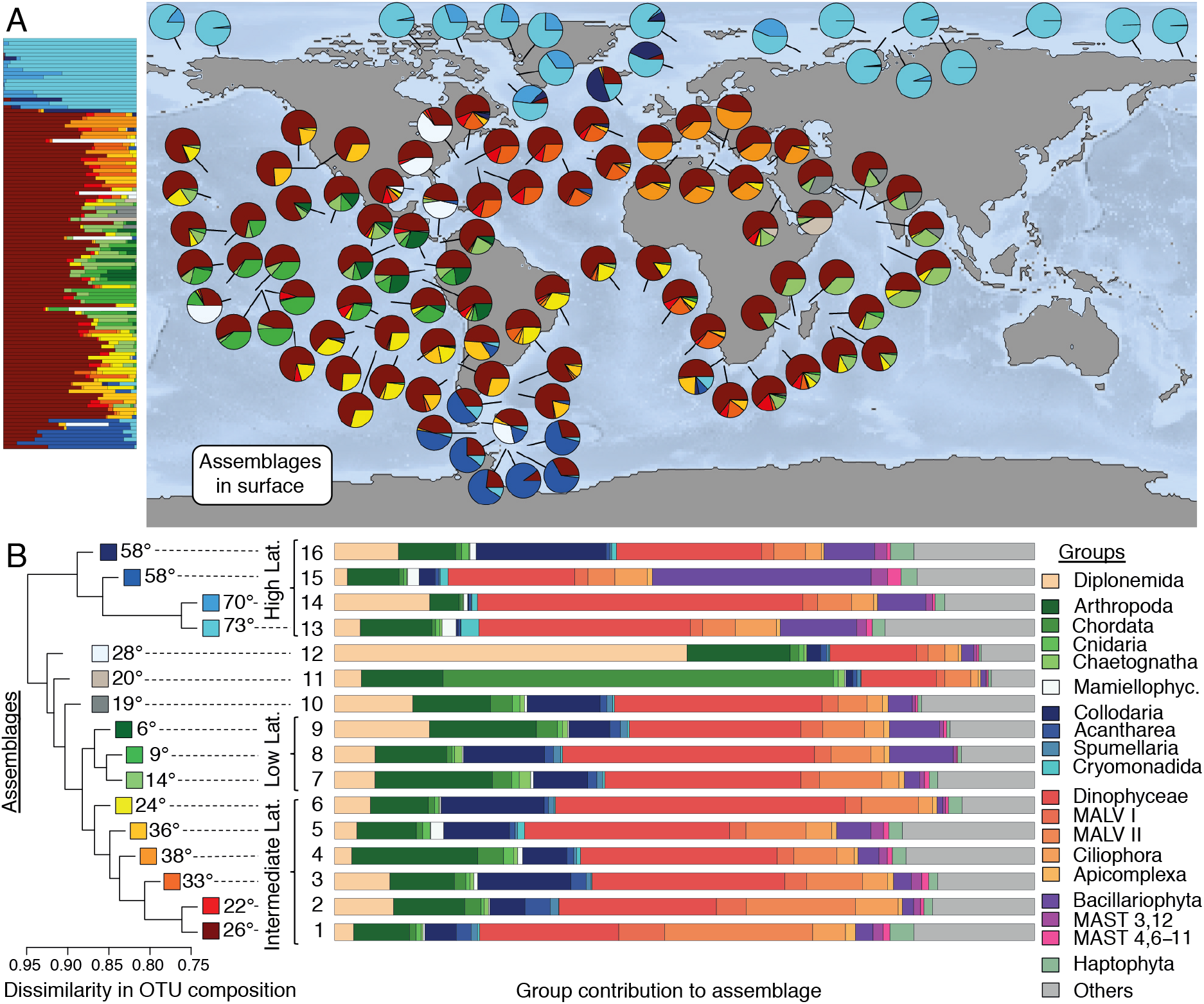
Global surface biogeography of eukaryotic plankton. The biogeography of all eukaryotic OTUs across *Tara* Oceans stations is characterized by 16 assemblages of co-occurring OTUs, each represented by a distinct color (in A and left side of B) and identified by a number from 1 to 16 (in B). (**A**) Relative contribution of the 16 assemblages to surface plankton community in *Tara* Oceans stations, represented as pies on the world map, and as stacked bars vertically ordered by latitude on the left-hand side of the map. (**B**) To the left: dendrogram of assemblage dissimilarity with respect to their composition in OTUs (Simpson dissimilarity). The mean absolute latitude at which each assemblage is found is indicated. Three clusters can be distinguished: a high-latitude cluster — the most distinctive — in shades of blue, an intermediate-latidude cluster in shades from yellow to red, and a low-latitude cluster in shades of green. To the right: barplot displaying the contribution of major eukaryotic groups (deep-branching monophyletic groups) to assemblages. The 19 groups shown in the barplot are those tallying more than 1,000 OTUs, grouped by phylogenetic relatedness.

This global analysis hides a strong heterogeneity across the 70 most diversified deep-branching groups of eukaryotic plankton (Table S1). Comparing the biogeography of these major groups using a normalized information-theoretic metric of dissimilarity (*23*; cf. Mat. & Meth.), we found high pairwise dissimilarity values (ranging between 0.64 and 0.97; Fig. S2). This heterogeneity can be decomposed into two main interpretable axes of variation (Fig. 2; cf. Mat. & Meth.). The first axis reflects the *amount* of biogeographic structure: group position on this axis is positively correlated to short-distance spatial autocorrelation (Pearson’s correlation coefficient *ρ* = 0.91 at the surface; Fig. S3A), which measures the tendency for close-by communities to be composed of the same assemblages (cf. Mat. & Meth.). Groups scoring low on this axis are characterized by strong local variation, or “patchiness”. The second axis reflects the *nature* of the biogeographic structure: group position on this axis is positively correlated to the scale of biogeographic organization, which we measured as the characteristic distance at which spatial autocorrelation vanishes (*ρ* = 0.53, *p* = 10^−6^ at the surface; Fig. S3B) and which ranges from ~7,000 to ~14,400 km across groups. Group position on the second axis is also positively correlated to within-basin autocorrelation (*ρ* = 0.56, *p* = 10^−7^ at the surface; Fig. S3C), which measures the tendency for communities from the same oceanic basin (e.g., North Atlantic, South Atlantic, Mediterranean, Southern Ocean) to be composed of the same assemblages, and negatively correlated with latitudinal autocorrelation (*ρ* = −0.49, *p* = 10^−5^ at the surface; S3D), which measures the tendency for communities at the same latitude on both sides of the Equator to be composed of the same assemblages (cf. Mat. & Meth.). Results are similar at the DCM, although less pronounced (Fig. S4). The 70 groups of eukaryotic plankton cover the full spectra of biogeographies (Fig. 2, Fig. S5, Table S1), from those with weak spatial organization, or high patchiness (i.e., scoring low on the first axis, such as Collodaria or Basidiomycota), to those organized at large spatial scale by oceanic basin (i.e., scoring high on both axes, such as Chordata or Arthropoda), and those organized at smaller spatial scale and according to latitude (i.e., scoring high on the first and low on the second axis, such as Mamiellophyceae, Haptophyta or MAST 3,12). These striking differences across planktonic groups suggest that accounting for their specificities is crucial to understanding their biogeography.

**Fig. 2.**
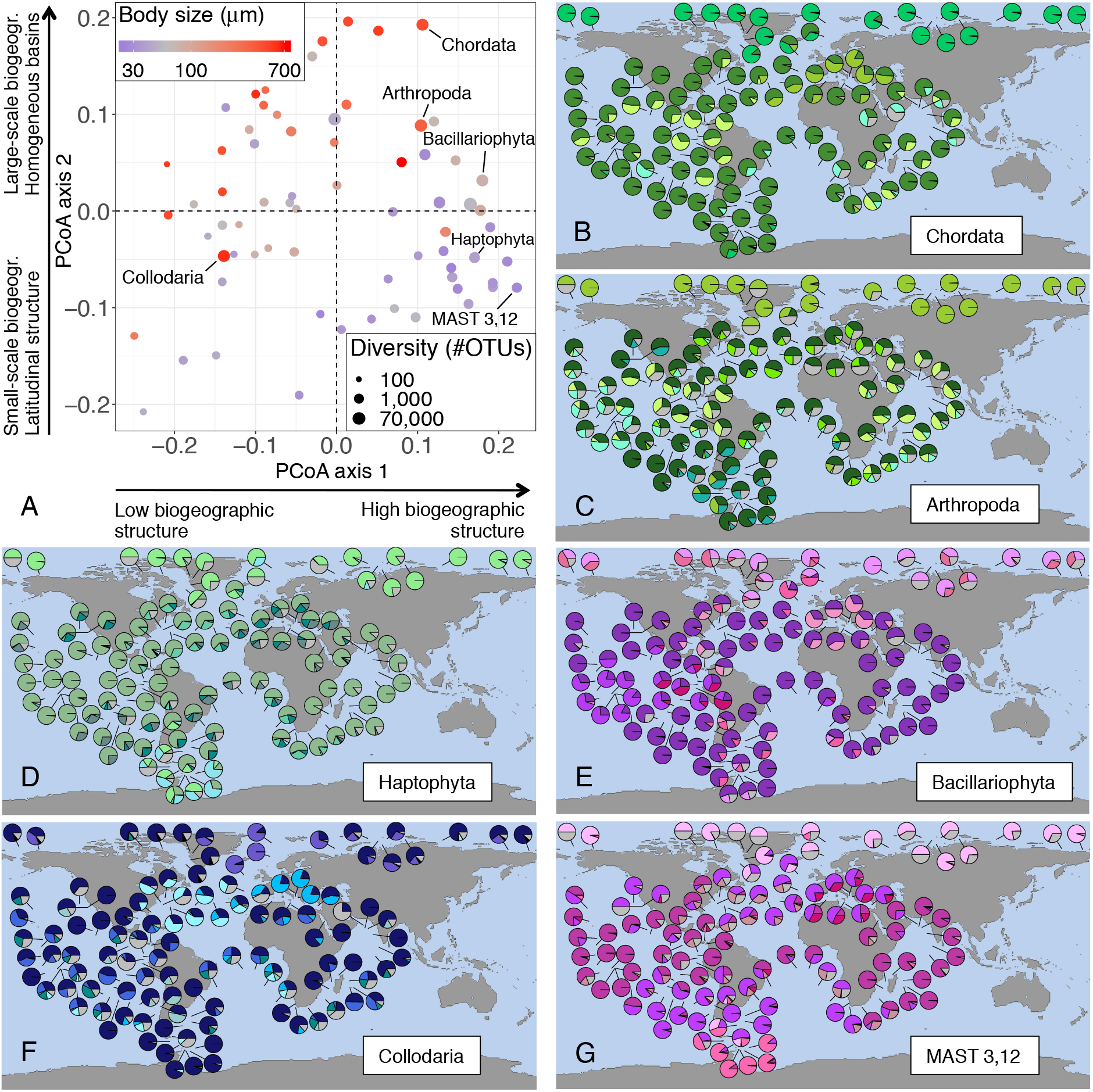
Biogeographic heterogeneity across major eukaryotic plankton groups. (**A**) Principal Coordinate Analysis (PCoA) of the biogeographic dissimilarity between 70 major groups of eukaryotic plankton. Each dot corresponds to the projection of a specific plankton group onto the first two axes of variation. Position along the first axis reflects the amount of biogeographic structure displayed by the group, from a patchy distribution with weak short-distance spatial autocorrelation on the left to a structured distribution with strong short-distance spatial autocorrelation on the right. Position along the second axis reflects the nature of biogeographic structure, from a biogeography structured by latitude at the bottom to a biogeography structured by oceanic basins at the top, as well as the scale of biogeographic organization, from small to large scale. Dot size is proportional to the log diversity of the corresponding group, and dot color represents its mean log body-size, from small (blue) to large (red). (**B**-**G**) Surface biogeography of six major eukaryotic plankton groups. The relative contribution of the 5 to 7 most prevalent assemblages is shown in color, and that of the remaining assemblages is shown in gray; the color used for the most prevalent assemblage corresponds to the color used in Fig. 1B for the corresponding group.

We investigated how biogeographic differences among major groups relate to their diversity, body size, and ecology, coarsely defined as either phototroph, phagotroph, metazoan or parasite (cf. Mat. & Meth.). We found that the amount of biogeographic structure (group position on the first axis) is strongly correlated to diversity (*ρ* = 0.77, *p* = 10^−13^ below 2,000 OTUs; Fig. 3A). This suggests that the maintenance of eukaryotic plankton diversity over ecological and possibly evolutionary scales is tightly linked to biogeographic structure, which may for example promote endemism. This relationship vanishes however for groups larger than about 2,000 OTUs, and two of the most diverse groups (Diplonemida, 38,769 OTUs and Collodaria, 17,417 OTUs) exhibit comparatively weak biogeographic structure. The amount of biogeographic structure is weakly anticorrelated to body size (*ρ* = −0.32, *p* = 0.007; Fig. S6A), and after accounting for differences in diversity across groups, is lower for metazoans than for phototrophs (ANCOVA t-test: *p* = 0.04, Fig. S6B), in agreement with the expectation of a higher local patchiness in their distribution induced by turbulent stirring (*15*, *24*). In contrast, the nature of biogeographic structure (group position on the second axis) is strongly correlated to body size (*ρ* = 0.64, *p* = 10^−9^ Fig. 3B) and ecology (ANOVA F-test: *p* = 10^−7^, Fig. 3C), and only weakly to diversity (*ρ* = 0.24, *p* = 0.05; Fig. S6C). Metazoan groups score high on the second axis of variation (with the notable exception of Porifera sponges, probably at the larval stage, which are excluded from statistical results) and phototrophs score low, while phagotrophs occupy an intermediate position, spanning a comparatively wider range of biogeographies (Fig. 3C). Parasites are just below metazoans, which suggests that their biogeography is influenced by that of their hosts. While body size covaries with ecology (phagotrophs are larger than phototrophs on average, and metazoans significantly larger than other plankton types; Fig. S7), the positive relationship between group position on the second axis and body size still holds within each of the four ecological categories (ANCOVA F-test: *p* = 10^−4^ Fig. S8). Diatoms (Bacillariophyta) are a striking example: of all phototrophs, they have the largest body size and also score highest on the second axis of variation. Conversely, ecology significantly influences group position on the second axis even after accounting for body size differences (ANCOVA F-test: *p* = 0.01). Collodaria, which we did not assign to an ecological category, score lower than expected from their large body size, but close to the average for phagotrophic groups (Fig. 2, Table S1). These results suggest that biogeographic patterns are influenced by both body size and ecology. To summarize, diversity-rich groups are biogeographically structured, with large-bodied heterotrophs (metazoans such as Copepods and Tunicates) exhibiting biogeographic variations at the scale of oceanic basins or larger, and small-bodied phototrophs (such as Haptophyta) at smaller spatial scale and following latitude (Fig. 2).

**Fig. 3.**
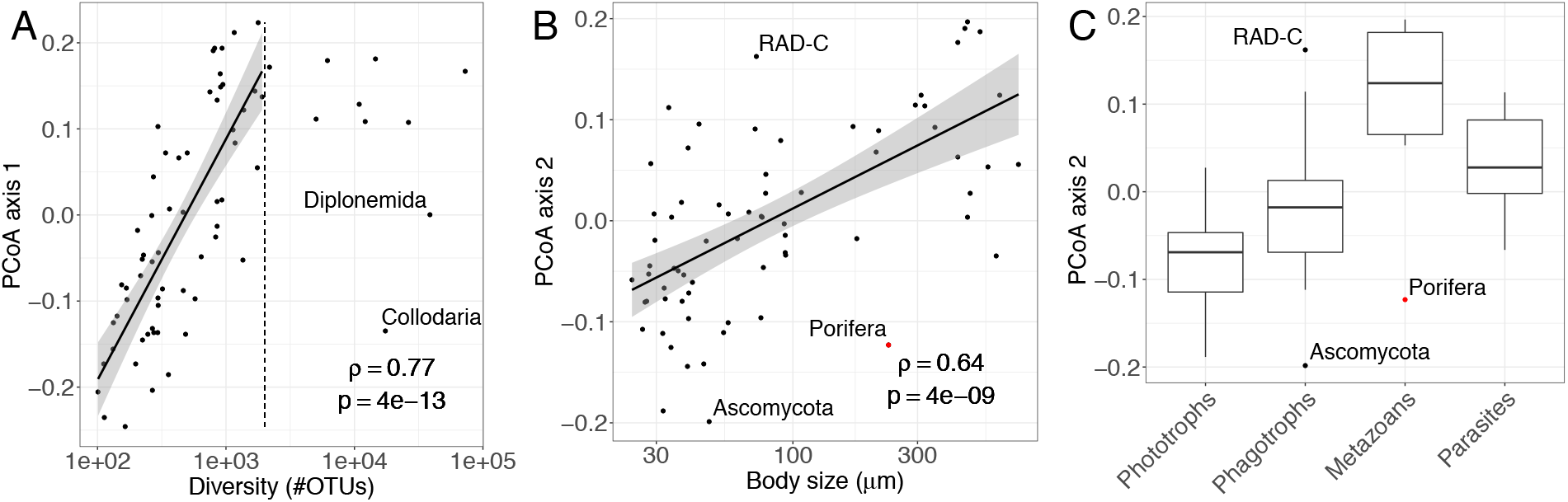
Relationship between biogeography and diversity, mean body size and ecology across major eukaryotic plankton groups. (**A**) The position of the 70 plankton groups along the first axis of biogeographic variation, indicative of the amount of biogeographic structure, increases sharply with log diversity (number of OTUs in the group) up to approximately 2,000 OTUs, but not beyond (as exemplified by Diplonemida and Collodaria, two of the most diverse groups). (**B**) The position of the 70 plankton groups along the second axis, indicative of the nature and spatial scale of biogeographic structure, increases with log mean body size, indicating that large-bodied plankton is organized at larger spatial scale and according to oceanic basins rather than latitude. (**C**) Positions along the second axis of plankton groups binned into four broad ecological categories (Collodaria and Dynophyceae were not categorized and are therefore not represented). Pairwise differences are all significant except between Phagotrophs and Parasites. The grey dot denotes Porifera, an outlier group excluded from statistical tests.

A global biogeography matching oceanic basins suggests that communities respond to environmental variations slowly enough to be homogenised by ocean circulation at the basin scale (i.e., gyres; *12*), but have little ability to disperse between basins, either due to the comparatively limited connectivity by currents or to environmental barriers, and therefore that their biogeography is primarily shaped by the main ocean currents (*13*). Conversely, a biogeography matching latitude, symmetric with respect to the Equator, suggests a faster response of communities to environmental variations within basins (which are structured by latitude and currents, e.g. the cross-latitudinal influence of the Gulf Stream), low cross-basin dispersal limitation, and therefore a comparatively more important role of local environmental filtering in shaping biogeography. To explain the global biogeography of major taxonomic groups, we compared biogeographic maps to maps of connectivity by currents and environmental conditions. We transformed the matrix of minimum transport times between pairs of stations, previously computed from a global ocean circulation model (*12*, *25*), into spatial patterns at different scales through eigenvector decomposition, thus obtaining a set of so-called Moran Eigenvector maps (thereafter simply referred to as “connectivity maps”; cf. Mat. & Meth.). These maps represent the hypothetical geographic patterns expected for plankton with temporal variation along currents matching these scales (Fig. S9, S10). We estimated local abiotic conditions using yearly-averaged measurements of temperature, nutrient concentration and oxygen availability (World Ocean Atlas 2013; *26*; cf. Mat. & Meth.). Because biotic interactions (predation, competition, parasitic and mutualistic symbiosis) are thought to be important determinants of plankton community structure (*27*), we also quantified local biotic conditions using the relative read counts of major eukaryotic groups (excluding the focal group; cf. Mat. & Meth.). Biotic conditions, similarly to abiotic ones, have a latitudinal structure, and we refer here to them collectively as ‘environmental conditions’ (Fig. S11, S12). The resulting environmental maps can be interpreted as the hypothetical geographic patterns expected for organisms with a fast response to local environmental conditions and whose dispersal by currents is not limiting. Hence, a biogeography matching connectivity maps better than environmental maps suggest that the constraints imposed by the seascape, that is the transport of plankton by oceanic currents modulated by mixing and ecological drift, but also by the responses to nutrient supplies and temperature variations during transport, dominate over those imposed by detectable local environmental filtering (see also *12*).

We found that the total variance in surface community composition that can be explained by connectivity maps and local environmental conditions (abiotic and biotic) averages 27% across groups (min. 0% for Porifera, max. 58%) and is, as expected, tightly correlated to the amount of biogeographic structure (*ρ* = 0.88; Fig. 4A; cf. Mat. & Meth.). The part of the variance that is statistically explained by connectivity patterns is primarily contributed by between-basin connectivity patterns (Fig. S10 & S13), and is for most groups larger than the part of the variance statistically explained by environmental data (at the surface, on average 40% of the explained variance is purely explained by connectivity versus 22% by the environment; Fig. S14A). This supports a prominent role of transport by the main current systems and of the processes occurring along those pathways in shaping eukaryotic plankton biogeography, both by extending the distribution of some taxa beyond their optimal range (*28*) and by constraining long-distance dispersal. Unmeasured environmental variations along currents likely contribute to this role of ocean circulation. As expected from our previous results, the ratio of the fractions of variance explained by connectivity patterns and environmental data, which reflects their relative contributions to biogeography, increases with group position on the second axis of variation (*ρ* = 0.44, *p* = 10^−4^; Fig. 4B). Accordingly, the relative contribution of connectivity by currents also increases with average group body size (*ρ* = 0.42, *p* = 10^−4^; Fig. 4C) and depends on ecology (ANOVA F-test: *p* = 0.003; Fig. 4D). These results indicate that metazoans are closer to drifting tracers strongly influenced by currents, and constrained in particular by limited between-basin connectivity, while phototrophs are more strongly coupled with environmental factors and disperse more readily between basins. The difference in sensitivity to local environmental conditions can be explained by differences in ecological requirements and community dynamics. Why there is a difference in between-basins dispersal is less clear. All basins are connected by currents within a few years of transport time (*29*), and small phototrophs may have a higher ability to disperse through environmental barriers by forming spores or dormant states (*10*). Alternatively, the looser environmental coupling and slower dynamics of metazoan communities might make them more sensitive to the smaller between-basin compared to within-basin water flow. Finally, within the variance explained by the local environment, an approximately equal share can be attributed to biotic and abiotic conditions for most groups (respectively 29% and 26% purely explained at the surface, on average; Fig. S14B), irrespective of their body size, ecology, diversity or biogeography (Fig. S15). Results are similar at the DCM, but are far less pronounced (Fig. S16, S17). Although we cannot exclude the possibility that local biotic conditions reflect the indirect effect of local abiotic factors that are not accounted for in our study, such as fluxes of nutrients, which are often more relevant to planktonic organisms than instantaneous nutrient concentrations (*28*), these results indicate an additional role for interspecific interactions in shaping community composition (*27*, *30*).

**Fig. 4.**
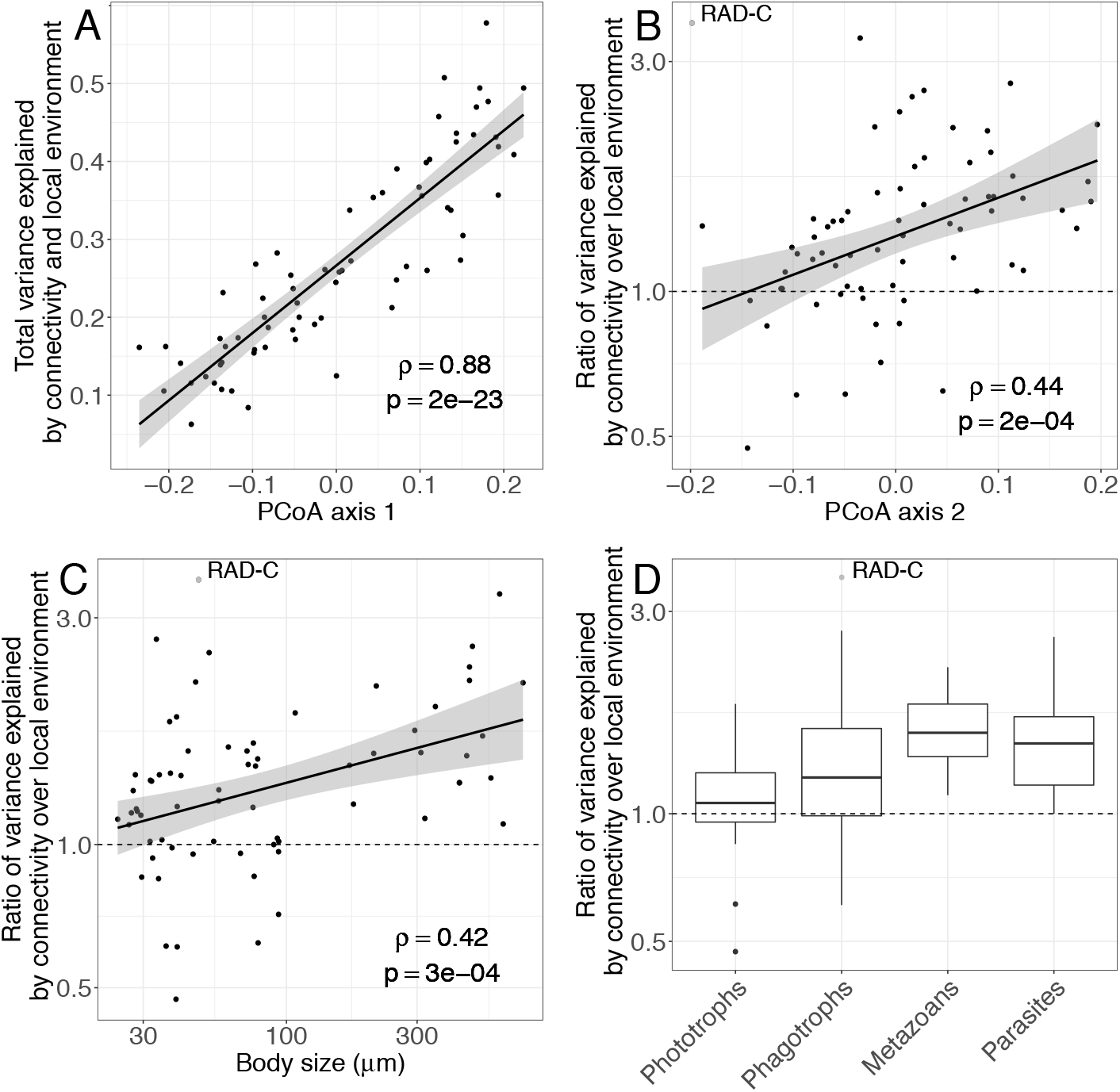
Drivers of surface biogeography across major eukaryotic plankton groups. (**A**) The total variance in surface biogeography that can be explained by the combination of connectivity by currents and (abiotic and biotic) local environmental conditions increases with the position of plankton groups on the first axis of biogeographic variation. (**B-D**) Across major plankton groups, the log ratio of the variance explained by connectivity over the variance explained by (abiotic and biotic) local environmental conditions (B) increases with group position on the second axis of variation, (C) increases with log mean body size, and (D) varies across broad ecological categories (pairwise differences are significant except Metazoans-Parasites and Phagotrophs-Phototrophs). The ratio is higher than 1 for most groups, reflecting an overall stronger influence of connectivity by currents compared to local environmental conditions on plankton biogeography at the surface. The grey dot denotes RAD-C, an outlier group excluded from statistical tests. We did not find any significant explanatory variable for Porifera and therefore excluded this group from these analyses.

Our study clarifies the patterns and processes underlying the global biogeography of the main groups of eukaryotic plankton in the sunlit ocean. Consistent with metagenomic results at lower taxonomic resolution (*12*), we find that eukaryotic plankton exhibits a global-scale biogeography, and that community variation is slow enough along currents to allow them to be the dominant driver of this biogeography. The continuous movement of water masses generates biogeographic patterns that are better represented by overlapping taxa assemblages than by the well-delineated biomes characteristic of terrestrial systems. Our comparison of eukaryotic plankton groups reveals several additional results. First, the geographic structuring induced by currents may have favored the generation and maintenance of eukaryotic plankton diversity. Second, plankton ecology matters beyond body size differences, and reciprocally body size matters beyond ecological differences. Third, body size and ecology influence primarily the nature of biogeographic patterns, namely their spatial scale of organization and whether they are organized by oceanic basins or latitude, and only secondarily the amount of biogeographic structure, namely local patchiness. Fourth, biotic conditions appear to be at least as important a driver of biogeography as local abiotic conditions. Our results reconcile the views that larger-bodied organisms are more dispersal-limited (*10*, *11*) and yet display a slower compositional turnover along currents than smaller organisms (*12*): at the global scale, organisms of larger sizes are indeed more dispersal-limited; however at the regional scale, they have wider spatial distributions, presumably linked to their specific ecologies, longer lifespan and reduced sensitivity to local environmental variations. At the two extremes, metazoan heterotrophs are structured at the scale of oceanic basins following the main currents, while small phototrophs are structured latitudinally with a comparatively larger influence of local environmental conditions, including biotic ones. Together, our results suggest that predictive modeling of plankton communities in a changing environment (*17*, *31*) will critically depend on our ability to model the impact of changes in ocean currents and to develop niche models accounting for both species ecology and interspecific interactions.

## Acknowledgements

We are grateful to Federico Ibarbalz for his essential help with the data. We thank Olivier Jaillon and Colomban de Vargas for feedback and early discusions on the project. We thank Mick Follows and Oliver Jahn for sharing MITgcm simulation results for the Arctic Ocean. We thank Florian Hartig and Odile Maliet for their guidance on Bayesian inference, Leandro Arístide and Felipe Delestro for their kind assistance with the figures, and Carmelo Fruciano and Benoît Perez for their advice on statistics. We thank Fabio Benedetti, Julien Clavel, Elena Kazamia, Sophia Lambert, Eric Lewitus, Marc Manceau, Olivier Missa, Silvia de Monte, Isaac Overcast, Ignacio Quintero, Enrico Ser-Giacomi, Ana Catarina Silva and Flora Vincent for suggestions and fruitful discussions.

## Author contributions

GSK and HM designed the study with the help of RW, DI and CB. GSK performed the analyses. RW contributed the transport time data and their interpretation. GSK and HM wrote the paper with substantial input from RW, DI and CB.

## Data availability

All data reported herein are available without restrictions. Metabarcoding data have been deposited at the European Nucleotide Archive (ENA) under accession numbers PRJEB6610 and PRJEB9737. Sample metadata are available from https://doi.org/10.1594/PANGAEA.875582.

## List of Supplementary Material

Materials and Methods

Appendix

Figures S1 to S17

Table S1

References (*32*-*47*)

## References and Notes

1. C. B. Field, M. J. Behrenfeld, J. T. Randerson, P. Falkowski, Primary production of the biosphere: integrating terrestrial and oceanic components. Science. 281, 237–240 (1998).

2. A. Z. Worden, M. J. Follows, S. J. Giovannoni, S. Wilken, A. E. Zimmerman, P. J. Keeling, Rethinking the marine carbon cycle: Factoring in the multifarious lifestyles of microbes. Science. 347, 1257594 (2015).

3. G. Beaugrand, R. R. Kirby, How Do Marine Pelagic Species Respond to Climate Change? Theories and Observations. Annual Review of Marine Science. 10, 169–197 (2018).

4. E. J. Raes, L. Bodrossy, J. van de Kamp, A. Bissett, M. Ostrowski, M. V. Brown, S. L. S. Sow, B. Sloyan, A. M. Waite, Oceanographic boundaries constrain microbial diversity gradients in the South Pacific Ocean. Proceedings of the National Academy of Sciences. 115, E8266–E8275 (2018).

5. D. Righetti, M. Vogt, N. Gruber, A. Psomas, N. E. Zimmermann, Global pattern of phytoplankton diversity driven by temperature and environmental variability. Science Advances. 5, eaau6253 (2019).

6. D. P. Tittensor, C. Mora, W. Jetz, H. K. Lotze, D. Ricard, E. V. Berghe, B. Worm, Global patterns and predictors of marine biodiversity across taxa. Nature. 466, 1098–1101 (2010).

7. M. T. Kavanaugh, M. J. Oliver, F. P. Chavez, R. M. Letelier, F. E. Muller-Karger, S. C. Doney, Seascapes as a new vernacular for pelagic ocean monitoring, management and conservation. ICES J Mar Sci. 73, 1839–1850 (2016).

8. M. Lévy, O. Jahn, S. Dutkiewicz, M. J. Follows, Phytoplankton diversity and community structure affected by oceanic dispersal and mesoscale turbulence. Limnol. Oceanogr. 4, 67–84 (2014).

9. C. de Vargas, S. Audic, N. Henry, J. Decelle, F. Mahe, R. Logares, E. Lara, C. Berney, N. Le Bescot, I. Probert, M. Carmichael, J. Poulain, S. Romac, S. Colin, J.-M. Aury, L. Bittner, S. Chaffron, M. Dunthorn, S. Engelen, O. Flegontova, L. Guidi, A. Horak, O. Jaillon, G. Lima-Mendez, J. Lukes, S. Malviya, R. Morard, M. Mulot, E. Scalco, R. Siano, F. Vincent, A. Zingone, C. Dimier, M. Picheral, S. Searson, S. Kandels-Lewis, S. G. Acinas, P. Bork, C. Bowler, G. Gorsky, N. Grimsley, P. Hingamp, D. Iudicone, F. Not, H. Ogata, S. Pesant, J. Raes, M. E. Sieracki, S. Speich, L. Stemmann, S. Sunagawa, J. Weissenbach, P. Wincker, E. Karsenti, C. Tara Oceans, Eukaryotic plankton diversity in the sunlit ocean. Science. 348, 1261605 (2015).

10. B. J. Finlay, Global Dispersal of Free-Living Microbial Eukaryote Species. Science. 296, 1061–1063 (2002).

11. E. Villarino, J. R. Watson, B. Jönsson, J. M. Gasol, G. Salazar, S. G. Acinas, M. Estrada, R. Massana, R. Logares, C. R. Giner, M. C. Pernice, M. P. Olivar, L. Citores, J. Corell, N. Rodríguez-Ezpeleta, J. L. Acuña, A. Molina-Ramírez, J. I. González-Gordillo, A. Cózar, E. Martí, J. A. Cuesta, S. Agustí, E. Fraile-Nuez, C. M. Duarte, X. Irigoien, G. Chust, Large-scale ocean connectivity and planktonic body size. Nature Communications. 9, 142 (2018).

12. D. J. Richter, R. Watteaux, T. Vannier, J. Leconte, P. Frémont, G. Reygondeau, N. Maillet, N. Henry, G. Benoit, A. Fernàndez-Guerra, S. Suweis, R. Narci, C. Berney, D. Eveillard, F. Gavory, L. Guidi, K. Labadie, E. Mahieu, J. Poulain, S. Romac, S. Roux, C. Dimier, S. Kandels, M. Picheral, S. Searson, T. O. Coordinators, S. Pesant, J.-M. Aury, J. R. Brum, C. Lemaitre, E. Pelletier, P. Bork, S. Sunagawa, L. Karp-Boss, C. Bowler, M. B. Sullivan, E. Karsenti, M. Mariadassou, I. Probert, P. Peterlongo, P. Wincker, C. de Vargas, M. R. d’Alcalà, D. Iudicone, O. Jaillon, T. O. Coordinators, Genomic evidence for global ocean plankton biogeography shaped by large-scale current systems. bioRxiv, 867739 (2019).

13. F. L. Hellweger, E. van Sebille, N. D. Fredrick, Biogeographic patterns in ocean microbes emerge in a neutral agent-based model. Science. 345, 1346–1349 (2014).

14. M.-A. Madoui, J. Poulain, K. Sugier, M. Wessner, B. Noel, L. Berline, K. Labadie, A. Cornils, L. Blanco-Bercial, L. Stemmann, J.-L. Jamet, P. Wincker, New insights into global biogeography, population structure and natural selection from the genome of the epipelagic copepod Oithona. Molecular Ecology. 26, 4467–4482 (2017).

15. E. R. Abraham, The generation of plankton patchiness by turbulent stirring. Nature. 391, 577–580 (1998).

16. L. Oziel, A. Baudena, M. Ardyna, P. Massicotte, A. Randelhoff, J.-B. Sallée, R. B. Ingvaldsen, E. Devred, M. Babin, Faster Atlantic currents drive poleward expansion of temperate phytoplankton in the Arctic Ocean. Nat Commun. 11, 1–8 (2020).

17. F. M. Ibarbalz, N. Henry, M. C. Brandão, S. Martini, G. Busseni, H. Byrne, L. P. Coelho, H. Endo, J. M. Gasol, A. C. Gregory, F. Mahé, J. Rigonato, M. Royo-Llonch, G. Salazar, I. Sanz-Sáez, E. Scalco, D. Soviadan, A. A. Zayed, A. Zingone, K. Labadie, J. Ferland, C. Marec, S. Kandels, M. Picheral, C. Dimier, J. Poulain, S. Pisarev, M. Carmichael, S. Pesant, S. G. Acinas, M. Babin, P. Bork, E. Boss, C. Bowler, G. Cochrane, C. de Vargas, M. Follows, G. Gorsky, N. Grimsley, L. Guidi, P. Hingamp, D. Iudicone, O. Jaillon, S. Kandels, L. Karp-Boss, E. Karsenti, F. Not, H. Ogata, S. Pesant, N. Poulton, J. Raes, C. Sardet, S. Speich, L. Stemmann, M. B. Sullivan, S. Sunagawa, P. Wincker, M. Babin, E. Boss, D. Iudicone, O. Jaillon, S. G. Acinas, H. Ogata, E. Pelletier, L. Stemmann, M. B. Sullivan, S. Sunagawa, L. Bopp, C. de Vargas, L. Karp-Boss, P. Wincker, F. Lombard, C. Bowler, L. Zinger, Global Trends in Marine Plankton Diversity across Kingdoms of Life. Cell. 179, 1084–1097.e21 (2019).

18. G. Sommeria-Klein, L. Zinger, E. Coissac, A. Iribar, H. Schimann, P. Taberlet, J. Chave, Latent Dirichlet Allocation reveals spatial and taxonomic structure in a DNA-based census of soil biodiversity from a tropical forest. Molecular Ecology Resources. 20, 371–386 (2019).

19. D. Valle, B. Baiser, C. W. Woodall, R. Chazdon, Decomposing biodiversity data using the Latent Dirichlet Allocation model, a probabilistic multivariate statistical method. Ecology Letters. 17, 1591–1601 (2014).

20. A. R. Wallace, The geographical distribution of animals: with a study of the relations of living and extinct faunas as elucidating the past changes of the earth’s surface (Cambridge University Press, 1876), vol. 1.

21. G. F. Ficetola, F. Mazel, W. Thuiller, Global determinants of zoogeographical boundaries. Nature Ecology and Evolution. 1, 1–7 (2017).

22. L. D. Talley, Descriptive physical oceanography: an introduction (Academic press, 2011).

23. M. Meila, Comparing clusterings—an information based distance. Journal of Multivariate Analysis. 98, 873–895 (2006).

24. A. Bertrand, D. Grados, F. Colas, S. Bertrand, X. Capet, A. Chaigneau, G. Vargas, A. Mousseigne, R. Fablet, Broad impacts of fine-scale dynamics on seascape structure from zooplankton to seabirds. Nat Commun. 5, 1–9 (2014).

25. S. Clayton, S. Dutkiewicz, O. Jahn, C. Hill, P. Heimbach, M. J. Follows, Biogeochemical versus ecological consequences of modeled ocean physics. Biogeosciences. 14, 2877–2889 (2017).

26. T. P. Boyer, J. I. Antonov, O. K. Baranova, C. Coleman, H. E. Garcia, A. Grodsky, D. R. Johnson, R. A. Locarnini, A. V. Mishonov, T. D. O’Brien, World Ocean Database 2013. (2013), doi:10.7289/V5NZ85MT.

27. G. Lima-Mendez, K. Faust, N. Henry, J. Decelle, S. Colin, F. Carcillo, S. Chaffron, J. C. Ignacio-Espinosa, S. Roux, F. Vincent, L. Bittner, Y. Darzi, J. Wang, S. Audic, L. Berline, G. Bontempi, A. M. Cabello, L. Coppola, F. M. Cornejo-Castillo, F. d’Ovidio, L. De Meester, I. Ferrera, M. J. Garet-Delmas, L. Guidi, E. Lara, S. Pesant, M. Royo-Llonch, G. Salazar, P. Sanchez, M. Sebastian, C. Souffreau, C. Dimier, M. Picheral, S. Searson, S. Kandels-Lewis, G. Gorsky, F. Not, H. Ogata, S. Speich, L. Stemmann, J. Weissenbach, P. Wincker, S. G. Acinas, S. Sunagawa, P. Bork, M. B. Sullivan, E. Karsenti, C. Bowler, C. de Vargas, J. Raes, C. Tara Oceans, Determinants of community structure in the global plankton interactome. Science. 348, 1262073 (2015).

28. S. Dutkiewicz, P. Cermeno, O. Jahn, M. J. Follows, A. E. Hickman, D. A. A. Taniguchi, B. A. Ward, Dimensions of marine phytoplankton diversity. Biogeosciences. 17, 609–634 (2020).

29. B. F. Jönsson, J. R. Watson, The timescales of global surface-ocean connectivity. Nature Communications. 7, 11239 (2016).

30. F. Vincent, C. Bowler, Diatoms Are Selective Segregators in Global Ocean Planktonic Communities. mSystems. 5(2020), doi:10.1128/mSystems.00444-19.

31. H. K. Lotze, D. P. Tittensor, A. Bryndum-Buchholz, T. D. Eddy, W. W. L. Cheung, E. D. Galbraith, M. Barange, N. Barrier, D. Bianchi, J. L. Blanchard, L. Bopp, M. Büchner, C. M. Bulman, D. A. Carozza, V. Christensen, M. Coll, J. P. Dunne, E. A. Fulton, S. Jennings, M. C. Jones, S. Mackinson, O. Maury, S. Niiranen, R. Oliveros-Ramos, T. Roy, J. A. Fernandes, J. Schewe, Y.-J. Shin, T. A. M. Silva, J. Steenbeek, C. A. Stock, P. Verley, J. Volkholz, N. D. Walker, B. Worm, Global ensemble projections reveal trophic amplification of ocean biomass declines with climate change. Proc. Natl. Acad. Sci. U.S.A. 116, 12907–12912 (2019).

32. F. Mahé, T. Rognes, C. Quince, C. de Vargas, M. Dunthorn, Swarm: robust and fast clustering method for amplicon-based studies. PeerJ. 2, e593 (2014).

33. L. Guillou, D. Bachar, S. Audic, D. Bass, C. Berney, L. Bittner, C. Boutte, G. Burgaud, C. de Vargas, J. Decelle, J. del Campo, J. R. Dolan, M. Dunthorn, B. Edvardsen, M. Holzmann, W. H. C. F. Kooistra, E. Lara, N. Le Bescot, R. Logares, F. Mahé, R. Massana, M. Montresor, R. Morard, F. Not, J. Pawlowski, I. Probert, A.-L. Sauvadet, R. Siano, T. Stoeck, D. Vaulot, P. Zimmermann, R. Christen, The Protist Ribosomal Reference database (PR2): a catalog of unicellular eukaryote Small Sub-Unit rRNA sequences with curated taxonomy. Nucleic Acids Res. 41, D597–D604 (2013).

34. D. M. Blei, A. Y. Ng, M. I. Jordan, Latent Dirichlet Allocation. Journal of Machine Learning Research. 3, 993–1022 (2003).

35. X.-H. Phan, L.-M. Nguyen, S. Horiguchi, in Proceeding of the 17th international conference on World Wide Web - WWW’08 (ACM Press, Beijing, China, 2008; http://portal.acm.org/citation.cfm?doid=1367497.1367510), p. 91.

36. B. Grün, K. Hornik, topicmodels: an R package for fitting topic models. Journal of statistical software. 40, 1–30 (2011).

37. R Core Team, R: A Language and Environment for Statistical Computing (R Foundation for Statistical Computing, Vienna, Austria, 2018).

38. E. Paradis, K. Schliep, ape 5.0: an environment for modern phylogenetics and evolutionary analyses in R. Bioinformatics. 35, 526–528 (2019).

39. P. Legendre, L. Legendre, Numerical Ecology (Elsevier, 2012).

40. D. Menemenlis, J.-M. Campin, P. Heimbach, C. Hill, T. Lee, M. Schodlok, H. Zhang, ECCO2: High Resolution Global Ocean and Sea Ice Data Synthesis. Mercator Ocean Quarterly Newsletter. 31, 13–21 (2008).

41. D. Chessel, A. Dufour, J. Thioulouse, The ade4 Package - I: One-Table Methods. R News, 5–10 (2004).

42. J. R. Watson, C. G. Hays, P. T. Raimondi, S. Mitarai, C. Dong, J. C. McWilliams, C. A. Blanchette, J. E. Caselle, D. A. Siegel, Currents connecting communities: nearshore community similarity and ocean circulation. Ecology. 92, 1193–1200 (2011).

43. D. Wilkins, E. van Sebille, S. R. Rintoul, F. M. Lauro, R. Cavicchioli, Advection shapes Southern Ocean microbial assemblages independent of distance and environment effects. Nature Communications. 4, 1–7 (2013).

44. J. Oksanen, F. G. Blanchet, M. Friendly, R. Kindt, P. Legendre, D. McGlinn, P. R. Minchin, R. B. O’Hara, G. L. Simpson, P. Solymos, M. H. H. Stevens, E. Szoecs, H. Wagner, vegan: Community Ecology Package (2018).

45. Clayton Sophie, Dutkiewicz Stephanie, Jahn Oliver, Follows Michael J., Dispersal, eddies, and the diversity of marine phytoplankton. Limnology and Oceanography: Fluids and Environments. 3, 182–197 (2012).

46. S. Dutkiewicz, P. Cermeno, O. Jahn, M. J. Follows, A. E. Hickman, D. A. A. Taniguchi, B. A. Ward, Dimensions of Marine Phytoplankton Diversity. Biogeosciences Discussions, 1–46 (2019).

47. J. Hausser, K. Strimmer, entropy: estimation of entropy, mutual information and related quantities (2014).

