## Supplementary Materials for "Global drivers of eukaryotic plankton biogeography in the sunlit ocean"

**This PDF file includes:**

Materials and Methods
Appendix
Figs. S1 to S17
Table S1

### Materials and Methods

#### DNA data processing

Planktonic organisms were sampled in 129 stations of the open ocean (no lagoon or costal waters) covering the Arctic, Atlantic, Indian, East Pacific and Southern Oceans as well as the Mediterranean and Red Seas. Samples were collected from subsurface mixed-layer waters (henceforth referred to as 'surface', about 5 m deep). In about half of the stations, samples were additionally collected at the Deep Chlorophyll Maximum ('DCM', ranging from 20 m to 190 m deep, most commonly around 40 m deep). At both depth levels, four different fractions of organisms' body size were collected: 0.8-5 mm, 5-20 mm (or 3-20 mm in some stations, which we treated as equivalent), 20-180 mm, and 180-2000 mm. In Arctic stations, a small size fraction without upper size limit (0.8 mm – infinity) was collected in place of the 0.8-5 mm size fraction. We treated both fractions as equivalent, since they were found to be of similar composition in stations where both were collected (indeed, small organisms greatly outnumber larger ones).

Whole DNA was extracted from these samples, then the V9 region of the gene coding for the eukaryotic 18S rRNA was PCR-amplified and the resulting amplicons were sequenced by Illumina sequencing. Sequencing reads were trimmed for quality, length and fidelity of primer sequences, then clustered into Operational Taxonomic Units (henceforth 'OTUs') using the SWARM unsupervised algorithm (*32*). OTUs were given taxonomic assignations by matching their most abundant sequence to a custom database derived from the Protist Ribosomal Reference (PR2; *33*). OTUs with less than 80% similarity to the closest reference sequence were discarded, as well as OTUs matching non-eukaryotic reference sequences. This pipeline resulted in a list of OTUs and their associated read count for each sample. See de Vargas et al. (*9*) for further detail on the sampling, wetlab and bioinformatics protocols. Taxonomic assignations of OTUs were then used to obtain ecological annotations based on literature, from which OTUs could be broadly classified into parasites, phototrophs, phagotrophs and metazoans (*17*).

For every station and depth, we pooled the results obtained for the four size fractions into a single aggregated sample (henceforth simply referred to as a 'sample'). We discarded the samples where one or more size fractions were missing so as not to bias the results. This treatment resulted in retaining 113 stations, broken down into 110 surface samples and 62 DCM samples and encompassing 250,057 OTUs.

#### Characterizing samples as mixtures of assemblages using Latent Dirichlet Allocation

To capture the spatial patterns of OTU co-occurrence across samples, we used a model-based algorithm of dimensionality reduction, Latent Dirichlet Allocation (LDA; *34*). We considered that an OTU occurs in a sample when it is represented by at least one sequence read, and we discarded read count information. The method consists in fitting a so-called mixed membership model to the list of OTU occurrences in each sample (i.e., the community matrix). Even though the model formally assumes that OTUs can be observed several times in each sample (i.e., it assumes discrete abundance data rather than presence-absence data), this does not impair model fitting and interpretation for presence-absence data (*18*). The model assumes that OTU occurrences are sampled from a mixture of several (unobserved) assemblages. Each assemblage represents a set of OTUs that tend to co-occur across samples. The fitting process consists in inferring the $K$ most likely assemblages from the data, where the number $K$ of assemblages is fixed beforehand. Assemblages are defined by their OTU composition, both in terms of OTU identity and relative prevalence. The relative prevalence of an OTU in an assemblage is proportional to its number of occurrences across the samples where the assemblage is present. Assemblages may share OTUs, and samples may contain a mixture of coexisting assemblages. As a consequence the model is able to capture spatial patterns despite the presence of many ubiquitous OTUs, a typical trait of microbial communities, and to accommodate gradual changes in taxonomic composition across space. The model is little influenced by OTUs of rare occurrence, since those OTUs contribute little co-occurrence information. Symmetric Dirichlet priors are put on the mixture of assemblages in samples and on the mixture of OTUs in assemblages, with respective control parameters $a$ and $d$.

We fitted the model to all samples simultaneously, making no distinction between surface and DCM samples. We used the Gibbs sampling algorithm of Phan et al. (*35*), wrapped in the R package 'topicmodels' (*36*), with control parameters $\alpha = 0.1$ and $\delta = 0.1$. Values of $a$ and $d$ lower than 1 favor low spatial overlap and few shared OTUs between assemblages, respectively. Model output is chiefly influenced by $d$: values of $d$ close to 1 or higher led to solutions where very few widely distributed assemblages shared the bulk of OTUs. These solutions were associated with lower predictive power on held-out data (as measured by perplexity; see next paragraph) and lower posterior probability compared to lower $d$ values. We ran the MCMC (Markov Chain Monte Carlo) chains for 3,000 iterations starting from random assemblages. After the first 2,000 iterations (burn-in), we recorded samples every 25 iterations for the last 1,000 iterations (i.e., 40 MCMC samples per chain). MCMC samples are sets of values for all the model's latent variables, which follow the model's posterior distribution given the data once the chain has converged. The associated likelihood values are computed as part of the algorithm. Among the 40 MCMC samples, we picked that with likelihood closest to the mean across samples, as a proxy for the set of latent variable values maximizing the posterior distribution.

We selected the optimal number $K$ of assemblages by cross-validation. We partitioned the data into random sets of 10 samples, and fitted the model on the data while successively holding out each 10-sample validation set. We then measured the predictive power of each fitted model on the corresponding validation set. We measured it using perplexity, a decreasing function of predictive power defined as the geometric mean of the likelihood across OTU occurrences (perplexity function in R package 'topicmodels'; *36*). We compared the mean perplexity across validation sets for $K$ between 2 and 35, and picked the minimum value after smoothing the curve with a 6-degree-of-freedom spline (function smooth.spline, R package 'stats'; *37*). For large datasets, the mean perplexity as a function of $K$ may enter a plateau after an initial decrease (Fig. S1). As a heuristic means to select the $K$ value corresponding to the onset of the plateau, we first fitted the model to the whole dataset for the $K$ value with minimum mean perplexity, and used the number of assemblages obtained after removing all the assemblages with a cumulative prevalence across the dataset of less than one sample. We then fitted the model again for the number of assemblages thus obtained.

Once we had selected the $K$ value, we ran 100 independent MCMC chains on the whole dataset from random initial conditions. To check for potential insufficient mixing along the chains, we measured the similarity in the spatial distribution of assemblages across the chains (Table S1), using the metric defined in Sommeria-Klein et al. (*18*). We picked the chain with posterior probability closest to the mean across chains for the final interpretation.

Comparing assemblages

Each assemblage is characterized by a list of OTUs and their relative prevalence. When running LDA on the whole eukaryotic data set, we measured the pairwise dissimilarity between assemblages as the Simpson dissimilarity of their composition in OTUs. We then built an UPGMA tree out of the dissimilarity matrix to obtain a hierarchical clustering of assemblages (function *agnes*, R package 'cluster').

Major eukaryotic groups

After having first considered all eukaryotic OTUs combined, we sought to compare biogeographic patterns across major groups of eukaryotic plankton. To this end, we classified OTUs into deep-branching monophyletic groups based on taxonomic assignations, as in de Vargas et al. (*9*), and we discarded those tallying less than 100 OTUs. We obtained 70 groups tallying between 101 to 72,769 OTUs (Dinophyceae), for a total of 241,020 OTUs.

We classified eukaryotic groups into four broad ecological categories based on the dominant ecology of their constituent OTUs: parasites, phototrophs, phagotrophs and metazoans. All groups fell entirely or mostly into one of these categories, except Dinophyceae (various ecological functions, including many mixotrophs) and Collodaria (mostly phagotrophic photohosts), which we did not classify and thus excluded from our statistical comparisons to ecology.

We estimated the mean body size of each group based on the distribution of the corresponding sequence reads over the four size fractions and across samples. Specifically, we computed the mean body size $\langle d_G \rangle$ of group $G$ across samples as:

$$\langle d_G \rangle = \frac{1}{S} \sum_{i=1}^{S} \frac{\sum_{f=1}^{4} \sum_{t \in G} p_{t,f,i} d_f}{\sum_{f=1}^{4} \sum_{t \in G} p_{t,f,i}}$$

where $S$ is the number of samples, $d_f$ the mid-range body size of fraction $f$ (i.e., respectively 2.9 mm, 12.5 mm, 100 mm, and 1,090 mm for the four size fractions), and $p_{t,f,s} = n_{t,f,i} / \sum_t n_{t,f,i}$ the relative abundance of OTU $t$ in fraction $f$ of sample $i$, as inferred from the number $n_{t,f,i}$ of sequence reads assigned to it. Groups' mean body size ranges from 24 mm (Cryptophyta) to 731 mm (Chaetognatha).

Groups diversity and body size are independent from each other ($p$ = 0.25), but variation in body size partly overlaps with ecological categories: all pairs of ecological categories have significantly distinct body size except parasites and phagotrophs (Fig. S7).

Amount of biogeographic structure

To quantify the amount of biogeographic structure exhibited by a planktonic group, we computed, separately for surface and DCM samples, the short-distance spatial autocorrelation $I_k$ in the global distribution of each assemblage $k$ across stations. We measured $I_k$ using Moran's index (function Moran.I, R package 'ape'; *38*), defined as:

$$I_k = \frac{S}{\sum_{i=1}^{S} \sum_{j=1}^{S} w_{ij}} \frac{\sum_{i=1}^{S} \sum_{j=1}^{S} w_{ij} \left(\theta_i^k - \langle \theta^k \rangle\right)\left(\theta_j^k - \langle \theta^k \rangle\right)}{\sum_{i=1}^{S} \left(\theta_i^k - \langle \theta^k \rangle\right)^2}$$

where $S$ is the number of stations, $\theta_i^k$ the proportion of assemblage $k$ in station $i$ (i.e., $\sum_{k=1}^{K} \theta_i^k = 1$), $\langle \theta^k \rangle = \sum_{i=1}^{S} \theta_i^k / S$ its mean over stations, and $w_{ij} = w\left(d_{ij}\right)$ is a weight function that decreases with the spatial distance $d_{ij}$ between stations $i$ and $j$. We defined the spatial distance between two stations as the shortest path between them that follows Earth's surface without crossing land (Dijkstra's algorithm; 12). We chose an inverse-square weight function satisfying $w(\max d_{ij}) = 0$ and $w(\min d_{ij}) = 1$:

$$w_{ij} = w(d_{ij}) = \frac{\left(\frac{\max d_{ij}}{d_{ij}}\right)^2 - 1}{\left(\frac{\max d_{ij}}{\min d_{ij}}\right)^2 - 1}$$

where $mind_{ij}$ is about 100 km and $maxd_{ij}$ 23,500 km. We then computed the overall short-distance spatial autocorrelation $I$ in the biogeography as the weighted mean of $I_k$ over assemblages, using the mean assemblage proportions $\langle\theta^k\rangle$ as weights, separately for the surface and the DCM:

$$I = \sum_{k=1}^{K} \langle\theta^k\rangle I_k$$

Scale of biogeographic organization

We quantified the scale of biogeographic organization as the characteristic distance at which spatial autocorrelation vanishes. We measured this distance in surface and at the DCM by computing Moran's I with a step weight function taking value $w_{ij} = 1$ if $d_{ij} < d$ and $w_{ij} = 0$ otherwise, and by varying $d$ linearly between $\min d_{ij}$ and $\max d_{ij}$ over 20 increments: $d^n = \min d_{ij} + n\left(\max d_{ij} - \min d_{ij}\right)/20$ for $n$ between 1 and 20. Moran's I decreases first linearly with spatial distance $d$ and then vanishes asymptotically. We smoothed the $I(d)$ curve with a 5-degree-of-freedom spline, and then performed a linear regression (function *lm*, R package 'stats') on its linear domain. We defined the characteristic distance at which spatial autocorrelation vanishes as the x-axis intercept of the linear regression (i.e., $-b/a$, where $a$ and $b$ are the slope and y-axis intercept, respectively).

Autocorrelation within oceanic basins

We measured the spatial autocorrelation within oceanic basins by computing Moran's I with a step weight function taking value $w_{ij} = 1$ when stations $i$ and $j$ belong to the same oceanic basin and $w_{ij} = 0$ otherwise, separately at the surface and the DCM. We defined as separate oceanic basins the Arctic Ocean, North Atlantic Ocean, South Atlantic Ocean, Mediterranean Sea, Red Sea, Indian Ocean, North Pacific Ocean, South Pacific Ocean and Southern Ocean. We expect a correlation between short-distance and within-basin spatial autocorrelation, since both are computed as Moran's I using different weight functions. To take this into account, we divided for each group the within-basin autocorrelation by the short-distance autocorrelation in statistical analyses.

Latitudinal autocorrelation

To measure whether the same assemblages tend occur at the same absolute latitude on both sides of the Equator, we computed, separately at the surface and the DCM, Moran's I with a weight function taking value $w_{ij} = e^{-(|l_i|-|l_j|)^2/\sigma^2}$ when $\mathrm{sign}(l_i) = -\mathrm{sign}(l_j)$ and $w_{ij} = 0$

otherwise, where $l_i$ is the latitude of station $i$ in degrees. We used $\sigma^2 = 25$, the value that maximized latitudinal autocorrelation in the surface biogeography of all eukaryotic OTUs combined. As for within-basin autocorrelation, we divided for each group the latitudinal autocorrelation by the short-distance autocorrelation in statistical analyses.

Comparing biogeography across groups

We applied our LDA decomposition pipeline (see above) separately to each of the major groups. To compare the resulting biogeography across groups, we computed a measure of biogeographic dissimilarity between pairs of groups. We used the relative mutual information between the spatial distribution of assemblages, an information theoretic quantity closely related to the Variation of Information (*23*) but normalized by total entropy so as to make it insensitive to differences in number of assemblages between groups.

We note $\theta_1 = \left(\theta_{1,i}^{k_1}\right)_{i\in[\![1,S]\!]}^{k_1\in[\![1,K_1]\!]}$ and $\theta_2 = \left(\theta_{2,i}^{k_2}\right)_{i\in[\![1,S]\!]}^{k_2\in[\![1,K_2]\!]}$ the spatial distribution over the $S$ stations of the respectively $K_1$ and $K_2$ assemblages in the biogeographies of groups 1 and 2, with $\sum_{k_1=1}^{K_1} \theta_{1,i}^{k_1} = 1$ and $\sum_{k_2=1}^{K_2} \theta_{2,i}^{k_2} = 1$ for every station $i$. We computed the entropy $H(\theta_j)$ and the mutual information $I(\theta_1, \theta_2)$ between $\theta_1$ and $\theta_2$ as:

$$H(\theta_j) = -\sum_{k_j=1}^{K_j} \langle\theta^{k_j}\rangle \log\langle\theta^{k_j}\rangle$$

$$I(\theta_1, \theta_2) = \sum_{\{k_1,k_2\}\in[\![1,K_1]\!]\times[\![1,K_2]\!]} \langle\theta_1^{k_1}\theta_2^{k_2}\rangle \log\frac{\langle\theta_1^{k_1}\theta_2^{k_2}\rangle}{\langle\theta_1^{k_1}\rangle\langle\theta_2^{k_2}\rangle}$$

where $\langle . \rangle$ stands for the mean over the $S$ stations. The relative mutual information between $\theta_1$ and $\theta_2$ is then defined as:

$$\tilde{I}(\theta_1, \theta_2) = \frac{I(\theta_1, \theta_2)}{H(\theta_1) + H(\theta_2) - I(\theta_1, \theta_2)}$$

The similarity index $\tilde{I}(\theta_1, \theta_2)$ varies between 0 and 1, and can be transformed into a dissimilarity index by taking $1 - \tilde{I}(\theta_1, \theta_2)$.

We performed a Principal Coordinate Analysis (function *pcoa.all*, *39*) on the $1 - \tilde{I}$ dissimilarity matrix between the 70 major groups, resulting in 69 PCoA axes. We performed multivariate linear regressions (function 'lm') of the projections of groups onto the PCoA axes against six explanatory variables: the amount of biogeographic structure, the scale of biogeographic organization, the within-basin autocorrelation, the latitudinal autocorrelation, the logarithm of group diversity and the logarithm of group body size. Each of these explanatory variables explained a significant part of the variance in the groups' projections onto all PCoA axes ($p < 10^{-3}$). When considering each PCoA axis separately, groups' projections onto the first two PCoA axes could be well predicted by the combination of these six explanatory variables ($R^2_{adj.} = 0.86$, $p = 10^{-25}$ for the first axis, $R^2_{adj.} = 0.69$, $p = 10^{-15}$for the second axis), while this was not the case for subsequent PCoA axes ($R^2_{adj.} < 0.17$, $p \gtrsim 10^{-2}$). Therefore the first two PCoA axes carry most of the interpretable biogeographic variation across groups, and as a consequence we focused on the ordination of the groups along those two axes.

Comparing the effect of body size, diversity and ecology

We assessed correlations between continuous variables using Pearson's correlation coefficient and the associated t-test (function *cor.test*). We tested the effect of ecology (with four factor levels: phototrophs, phagotrophs, metazoans and parasites) on a continuous variable (i.e., group position on the first two PCoA axes, or a ratio of explained variances) by an Analysis of Variance (ANOVA), and the respective effects of ecology and a continuous covariate (either log body size or log diversity) by an Analysis of Covariance (ANCOVA; functions *lm* and *anova*). We considered the t-tests between pairs of ecological categories only when the F-test was significant, and grouped ecological categories together when this improved the model. We removed obvious outlier groups from statistical analyses, so as not to break the normality and heteroscedasticity assumptions (Porifera for group position on the second PCoA axis; RAD-C for the ratio of the fractions of variance explained by connectivity and the environment — see below). Including these outliers in the analyses did not qualitatively change the results. We used a 5% significance threshold.

Abiotic environmental variables

For each sample, we used as local abiotic conditions the mean annual values measured at the approximate location and depth of the sample for temperature, nitrate, phosphate and silicate concentrations, dissolved oxygen concentration, oxygen saturation and apparent oxygen utilization (World Ocean Atlas 2013; *26*). We also used iron concentration values derived from model simulations (*40*). We conducted a Principal Component Analysis (PCA) on these abiotic environmental variables, separately for surface and DCM samples, after centering and standardization (function dudi.pca, R package 'ade4'; *40*). We retained the first three axes for further analysis (axes with eigenvalue larger than 0.8).

For surface samples, the first axis amounts to 44% of the total variance (eigenvalue = 3.5), and corresponds to variation in temperature as well as in nitrate, phosphate, silicate and dissolved oxygen concentrations. The second axis amounts to 26% of variance (eigenvalue = 2.1) and corresponds to variation in oxygen saturation and utilization. The third axis amounts to 16% of variance (eigenvalue = 1.3) and is mostly driven by iron concentration (Fig. S11).

For DCM samples, the first axis amounts to 51% of the total variance (eigenvalue = 4.1), and corresponds mostly to variation in phosphate and nitrate concentration, as well as oxygen utilization and saturation. The second axis amounts to 27% of variance (eigenvalue = 2.2), and corresponds mostly to variation in temperature and dissolved oxygen concentration. The third axis amounts to 10% of variance (eigenvalue = 0.84) and is driven by iron concentration.

Biotic environmental variables

We used the relative abundances in the community of the 70 major groups of eukaryotic plankton under study as proxy for local biotic conditions. We estimated the local relative abundance $a_{G,i}$ of a group in sample $i$ as the mean of its relative read count in the four size fractions:

$$a_{G,i} = \frac{\sum_{f=1}^{4} \sum_{t \in G} p_{t,f,i}}{\sum_{f=1}^{4} \sum_{t} p_{t,f,i}}$$

where, as defined previously for the calculation of body size, $p_{t,f,i}$ is the relative read count of OTU $t$ in fraction $f$ of sample $i$. The quantity $a_{G,i}$ is not directly a measure of the relative number of individuals in group $G$, because it is obtained by summing over size fractions, and both the density of individuals per volume of water and the sampled volume of water differ widely among size fractions. It can nevertheless be used to characterize the variation in community composition across stations.

We conducted a Principal Component Analysis (PCA) on relative abundances $a_G$ across groups, separately for surface and DCM samples, after centring and standardization (function dudi.pca, R package 'ade4'; *40*), and we retained the axes with eigenvalue larger than 0.8 as biotic environmental variables for further analysis (the first 28 axes for surface samples; the first 23 axes for DCM samples; Fig. S12). To avoid using the abundance of the group under study as an explanatory variable, we performed 70 separate PCAs, each time removing the focal group.

Connectivity maps

To quantify the role of transport by currents in generating the observed biogeographies, we compared them with connectivity maps known as Moran Eigenvector Maps (MEMs), obtained by decomposing the matrix of pairwise minimum transport times between stations using Principal Coordinate Analysis (PCoA), as described below (*39*). In terrestrial ecology, similar maps are obtained by decomposing the matrix of pairwise geographic distances between sampled sites, and are classically used to assess the effect of dispersal limitation by distance on the distribution of species.

Here, we measure the connectivity of stations using minimum transport times between stations, in line with previous studies using Lagrangian transit times to explain the spatial distribution of marine plankton (*29*, *42*, *43*). This measure of connectivity is more robust than physical connectivity (i.e., the number of particles exchanged between stations), which strongly depends on the number of particles considered in the simulation as well as on the method used to reconstruct the trajectories of particles between stations. When seeking to explain patterns of taxon presence-absence for planktonic organisms, the minimum transport time between stations appears more relevant than the mean transport time, since only a few individuals are required to 'seed' a location with a given taxon (*29*, *43*). Moreover, mean transport times are not well-defined in the global ocean in the absence of a physically motivated upper time-scale (*29*). Finally, minimum transport time has been shown to be a good predictor of the average amount of change in global plankton community composition that takes place along currents over a timescale of a year (i.e. a few thousands km), as a result of mixing, environmental variations, internal biotic interactions, behaviour and random compositional drift (*12*).

The minimum transport times were computed by Richter et al. (*12*) using a numerical simulation of a global oceanic circulation model (MITgcm Darwin; *25*), as summarized here. In this simulation, particles were released uniformly across the globe and advected for a cycle of 6 years using the horizontal velocity field along with a turbulent diffusivity. A set of 10,000-year trajectories was then constructed using this 6-year master cycle with particles seeded in each sampling station. Transport times between sampled locations were inferred by considering every event when a particle travelled from one sampled location to another, up to a radius of 200 km (see *12* for more details). Only stations that had exchanged at least 10 particles were considered significantly connected. This computation was performed twice using simulations at 5-m depth and 75-m depth, so as to estimate the minimum transport times at the surface and at the DCM, respectively. We thus obtained two symmetric square matrices, one for surface samples and one for DCM samples, with minimum transport times as entries for connected pairs of stations and missing values for unconnected pairs.

From these two matrices of pairwise minimum transport times, we generated connectivity maps (MEMs) taking one value per station as follows (*39*). We first computed for each matrix a minimum spanning tree among samples using function *spantree* of R package 'vegan' (*44*). Following the recommendations of Legendre & Legendre (*39*), we truncated the matrix of minimum transport times to retain only those connections necessary to connect all stations together (i.e., to obtain a connex graph), if possible. For surface samples, we found that a single tree connected all stations as long as we retained all minimum transport times below 2.1 years (which corresponds to distances up to a few thousands km, cf. Fig. S9). By doing so, we effectively restricted ourselves to the range of minimum transport times over which minimum transport time increases approximately linearly with the geographic distance between stations. For DCM samples, no single spanning tree connected all stations, and so we chose to retain all minimum transport times below 3.15 years, which led to the Mediterranean, the Red Sea and the Southern Ocean being disconnected from the remaining samples. In both matrices, we set the diagonals and all the elements above the selected threshold to four times the threshold value, and we conducted a PCoA of the resulting truncated connectivity matrices (function *pcoa.all*, *39*). We obtained 61 eigenvectors associated with strictly positive eigenvalues for the surface connectivity matrix and 35 for the DCM connectivity matrix, which we used as connectivity maps at the surface and the DCM.

The resulting connectivity maps display patterns of connectivity at temporal and spatial scales ranging from a few days and a hundred km (the minimal distance between a pair of stations) up to the global scale, and can therefore be used to assess the influence of transport by currents both within and between ocean basins (Fig. S10), which is difficult to achieve when directly using pairwise transport times between stations. They identify oceanographic features that are known to support high connectivity, such as the North Atlantic gyre system, the eastward flow between Scandinavia and Siberia in the Arctic Ocean, the South Pacific gyre, the Mediterranean Sea cyclonic circulation and the western Indian Ocean gyre system (Fig. S10).

### Variation partitioning

To assess the influence of explanatory variables on biogeography, we compared their distribution across stations to that of assemblages through multivariate linear regression, after centering and standardization. We used the adjusted coefficient of multiple determination $R_a^2$ as a measure of the variance in the distribution of assemblages across stations (i.e., in the biogeography) that can be explained by a set of explanatory variables (function rda, R package 'vegan'; *44*). We considered three sets of explanatory variables: abiotic environmental variables, biotic environmental variables, and connectivity maps (see above). For each taxonomic group and within each set of variables, we tested whether each variable individually explained a significant amount of variance in the biogeography (functions *rda* and *anova*), separately for the surface and DCM sets of samples. We only retained in further analyses the variables that were significant at 5% after Benjamini-Hochberg correction for multiple comparison. At least one explanatory variable was retained for every group except Porifera, which were therefore excluded from the variance partitioning.

We partitioned the variance explained by two sets of variables $A$ and $B$, denoted by $R^2_{a,A\cap B}$, into the variance explained purely by $A$ and $B$, denoted by $\tilde{R}^2_{a,A}$ and $\tilde{R}^2_{a,B}$, and that explained jointly by $A$ and $B$, denoted by $\tilde{R}^2_{a,A\cap B}$:

$$R^2_{a,A\cap B} = \tilde{R}^2_{a,A} + \tilde{R}^2_{a,B} + \tilde{R}^2_{a,A\cap B}$$

This partitioning can be obtained from the variance independently explained by $A$ and $B$, denoted by $R^2_{a,A}$ and $R^2_{a,B}$, as follows (function *varpart*, R package ‘vegan’):

$$\tilde{R}^2_{a,A\cap B} = R^2_{a,A} + R^2_{a,B} - R^2_{a,A\cap B}$$
$$\tilde{R}^2_{a,A} = R^2_{a,A\cap B} - R^2_{a,B}$$
$$\tilde{R}^2_{a,B} = R^2_{a,A\cap B} - R^2_{a,A}$$

For each taxonomic group, we partitioned the variance explained by all retained explanatory variables into the variance purely explained by connectivity maps, that purely explained by environmental variables — lumping biotic and abiotic variables together, and the variance jointly explained by both sets of variables. We compared across taxonomic groups the following quantities: the total explained variance, the fraction of it purely explained by connectivity maps, the fraction of it purely explained by the local environment, and the ratio of the variance explained by connectivity maps (both purely and jointly) over that explained by the local environment (both purely and jointly).

We similarly partitioned the variance explained by all environmental variables into the variance purely explained by abiotic variables, that purely explained by biotic variables, and the variance jointly explained by both sets of variables. We then compared across taxonomic groups the fraction of the environmentally explained variance purely explained by biotic variables, that purely explained by abiotic variables, and the ratio of the variance explained by biotic variables (both purely and jointly) over that explained by abiotic variables (both purely and jointly).

### Appendix: biogeographic patterns for all eukaryotic OTUs combined

In this appendix, we investigate in more details the biogeographic patterns obtained by applying Latent Dirichlet Allocation to all eukaryotic OTUs combined.

#### Fine-scale global biogeography

Beyond the broad geographic pattern formed by the three dominant assemblages (cf. main text), a finer geographic structure emerges from the non-dominant assemblages (Fig. A1A&C). Part of this structure follows a latitudinal trend, with several assemblages (e.g. assemblage 3 in dark orange, 5 in dark yellow and 6 in bright yellow) occurring in geographically discontinuous zones at similar latitudes on both sides of the Equator. Besides latitude, the geographic distribution of planktonic communities appears to be influenced by oceanic basins, currents, and upwellings. For example, some assemblages are geographically restricted to particular oceanic basins: assemblages 4 (light orange; Mediterranean Sea), 10 (dark grey; Persian Gulf), 11 (light grey; Red Sea), 8 (light green; Central Equatorial Pacific), 9 (dark green; Eastern Equatorial Pacific) and 7 (bright green; Indian Ocean and Equatorial Pacific). Communities along the Gulf Stream are characterized by assemblage 3, and those along the Benguela, South Equatorial and Brazilian currents smoothly transition from assemblage 3 to 6 and then 5. Assemblage 5 (rich in phototrophic OTUs) is associated with temperate upwelling areas around the world. Finally, areas off the South African, South-Brazilian, South-Chilean and Ecuadorian coasts display the most complex admixtures of assemblages, with as many as nine assemblages represented in the three *Tara* Oceans stations off South Africa (Fig. 2A in the main text). These regions correspond to the diversity hot spots predicted by models (*45*, *46*).

As we did for specific clades (cf. Mat. & Meth.), this biogeographic pattern can be summarized by its amount of short-distance spatial autocorrelation, as well as by the characteristic distance at which spatial autocorrelation vanishes, which measures the scale of spatial organization. Spatial autocorrelation is strong at short distance (Moran's I = 0.75 at the surface; I = 0.62 at the DCM, Fig A1B), meaning that close-by stations tend to be composed of the same assemblages. Short-distance spatial autocorrelation is higher at the surface than at the DCM, which may reflect faster mixing by currents, but also possibly the higher variability in the depth at which DCM samples were measured compared to surface samples. Spatial autocorrelation then decreases with distance and vanishes to zero with a characteristic scale of about 11,900 km in surface and 11,200 km at the DCM, which is the approximate size of an oceanic basin (Fig. A1B).

#### Comparing surface and DCM samples

Planktonic communities appear much more structured by geography than by depth (Fig. A1A&C). Indeed, biogeographic patterns are largely similar between surface and DCM. All assemblages are represented at both the surface and the DCM, and only two of them are mostly characteristic of either the surface (assemblage 6 in bright yellow; rich in dinoflagellates) or the DCM (assemblage 2 in red; similar to assemblage 1 in its composition in major clades). Both are found in tropical waters, reflecting the weaker mixing of water layers at lower latitude. In subtropical waters, DCM-associated assemblage 2 becomes more present at the surface, reflecting the increased mixing of water layers at higher latitude.

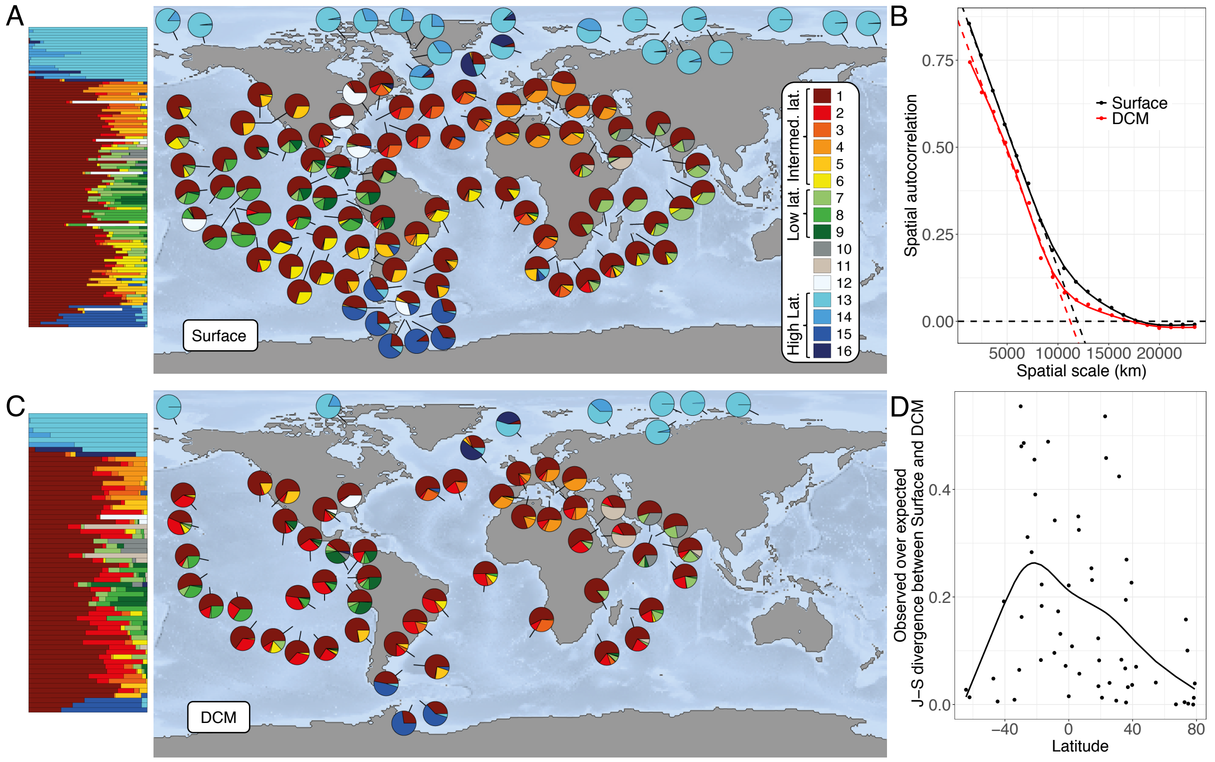


**Figure A1. Biogeography of eukaryotic plankton in surface and at the Deep Chlorophyll Maximum (DCM).** (**A**) Same as Figure 1A in the main text: relative contribution of assemblages to Tara stations at the surface, represented as pies on the world map, and also as stacked bars (vertically ordered by latitude) on the left-hand side of the map. (**B**) Spatial autocorrelation in the assemblage proportions as a function of the spatial scale considered, at the surface (black) and at the DCM (red). We define the characteristic scale of spatial autocorrelation as the intercept of the linear regression on the x-axis, which yields 11,900 km at the surface and 11,200 km at the DCM. (**C**) Relative contribution of biomes to Tara stations at the DCM. (**D**) Dissimilarity in biome composition between the surface and the DCM across Tara stations, as a function of latitude. Dissimilarity is computed as the Jensen-Shannon divergence between surface and DCM samples within a station, divided by the mean dissimilarity between surface and DCM across stations. Dissimilarity is much (about 6 times) lower within a station that across stations, and is lower at higher latitude.

To quantify the amount of similarity between surface and DCM samples in terms of assemblage composition, we computed the Jensen-Shannon divergence between the surface assemblage composition $\boldsymbol{\theta}_{Sur,i} = \left(\theta_{Sur,i}^{k}\right)^{k=[\![1,K]\!]}$ and the DCM assemblage composition $\boldsymbol{\theta}_{DCM,i} = \left(\theta_{DCM,i}^{k}\right)^{k=[\![1,K]\!]}$ for each station where samples were collected both in surface and at the DCM. Jensen-Shannon divergence is defined as:

$$\mathrm{JS}\left(\boldsymbol{\theta}_{Sur,i} \parallel \boldsymbol{\theta}_{DCM,i}\right) = \frac{1}{2}\mathrm{KL}\left(\boldsymbol{\theta}_{Sur,i} \parallel \frac{1}{2}\left(\boldsymbol{\theta}_{Sur,i} + \boldsymbol{\theta}_{DCM,i}\right)\right)$$

$$+\frac{1}{2}\mathrm{KL}\left(\boldsymbol{\theta}_{DCM,i} \parallel \frac{1}{2}\left(\boldsymbol{\theta}_{Sur,i} + \boldsymbol{\theta}_{DCM,i}\right)\right)$$

where $\mathrm{KL}\left(\boldsymbol{\theta}_i \parallel \boldsymbol{\theta}_j\right) = \sum_{k=1}^{K} \theta_i^k \log\left(\theta_i^k / \theta_j^k\right)$ is the Kullback-Leibler divergence between distributions $\boldsymbol{\theta}_i$ and $\boldsymbol{\theta}_j$ (function KL.plugin, R package ‘entropy’; *47*). We divided it by the mean Jensen-Shannon divergence across all pairs of surface and DCM samples, so as to assess whether surface and DCM samples of the same station had more similar decompositions than expected based on the overall similarity between the surface and DCM layers. We found that the composition in assemblages is on average 6 times more similar between a surface and a DCM sample if they come from the same station compared to a random pair of stations (Fig A1D). This similarity increases with latitude as the water column becomes less stratified.

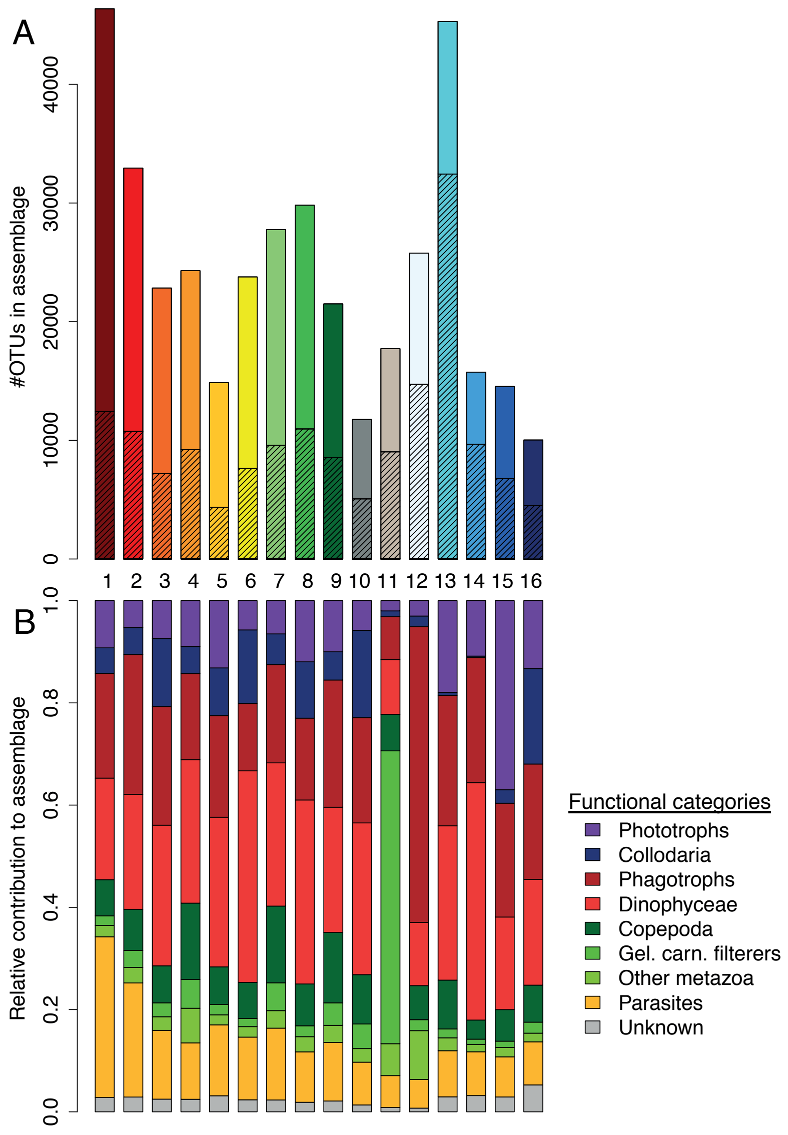


**Figure A2**: **OTU richness and functional composition of the 16 assemblages obtained for all Eukaryotic OTUs.** (**A**) Number of OTUs per assemblage, where asssemblages are identified by the numbers and colors used in Fig. 1 (main text) and Fig. A1. The hatched fraction represents for each assemblage the proportion of OTUs that are found only in that assemblage. No OTU is found in all assemblages. (**B**) Relative contribution of broad functional categories to each assemblage. Collodaria (mostly phagotrophic photohosts; in dark blue) and Dinophyceae (mostly mixotrophic; in light red) are not assigned to an ecological category and are represented separately. Copepods and gelatinous carnivore-filterers (i.e., Tunicates, Chaetognatha, Cnidaria and Ctenophora) are shown separately within metazoans.

### OTU composition of assemblages

The OTU composition is markedly distinct between assemblages (mean Simpson pairwise dissimilarity of 0.91). Each assemblage contains, on average, a significant proportion of OTUs that are not unique to that assemblage (61%), but half of these OTUs are shared by only two assemblages, and none by all (Fig. A2A). Assemblages nonetheless tend to have broadly similar compositions at the level of major planktonic clades (cf. Figure 1B in the main text). This is also true of their functional compositions (Fig. A2B). The only outliers are the Chordata-rich assemblage 11 characteristic of the Red Sea (mostly composed of

Tunicates), in light grey, and the diplonemid-rich assemblage 12, in white (Fig. 1B, Fig. A2B), which occurs in scattered locations.

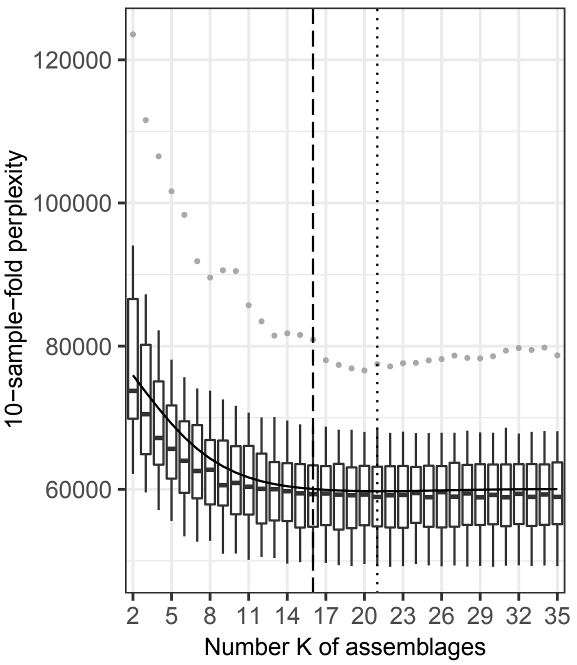


**Fig. S1. Selection of the number of assemblages *K* = 16 for all eukaryotic OTUs by cross-validation.** For each value of *K*, we randomly divided the samples into 10-sample folds; then for each fold, we fitted the model on the full data set minus that fold and we evaluated the perplexity of the resulting model on the fold. This plot shows the perplexity values obtained as a function of *K* (one point per held-out fold). The solid line indicates the mean perplexity across held-out folds per *K* value, and the dotted line indicates the *K* value for which mean perplexity reaches its minimum (*K* = 21). The dashed line indicates the selected *K* value (*K* = 16), obtained by fitting the model for *K* = 21 and removing all the assemblages with a cumulative prevalence across samples of less than one sample.

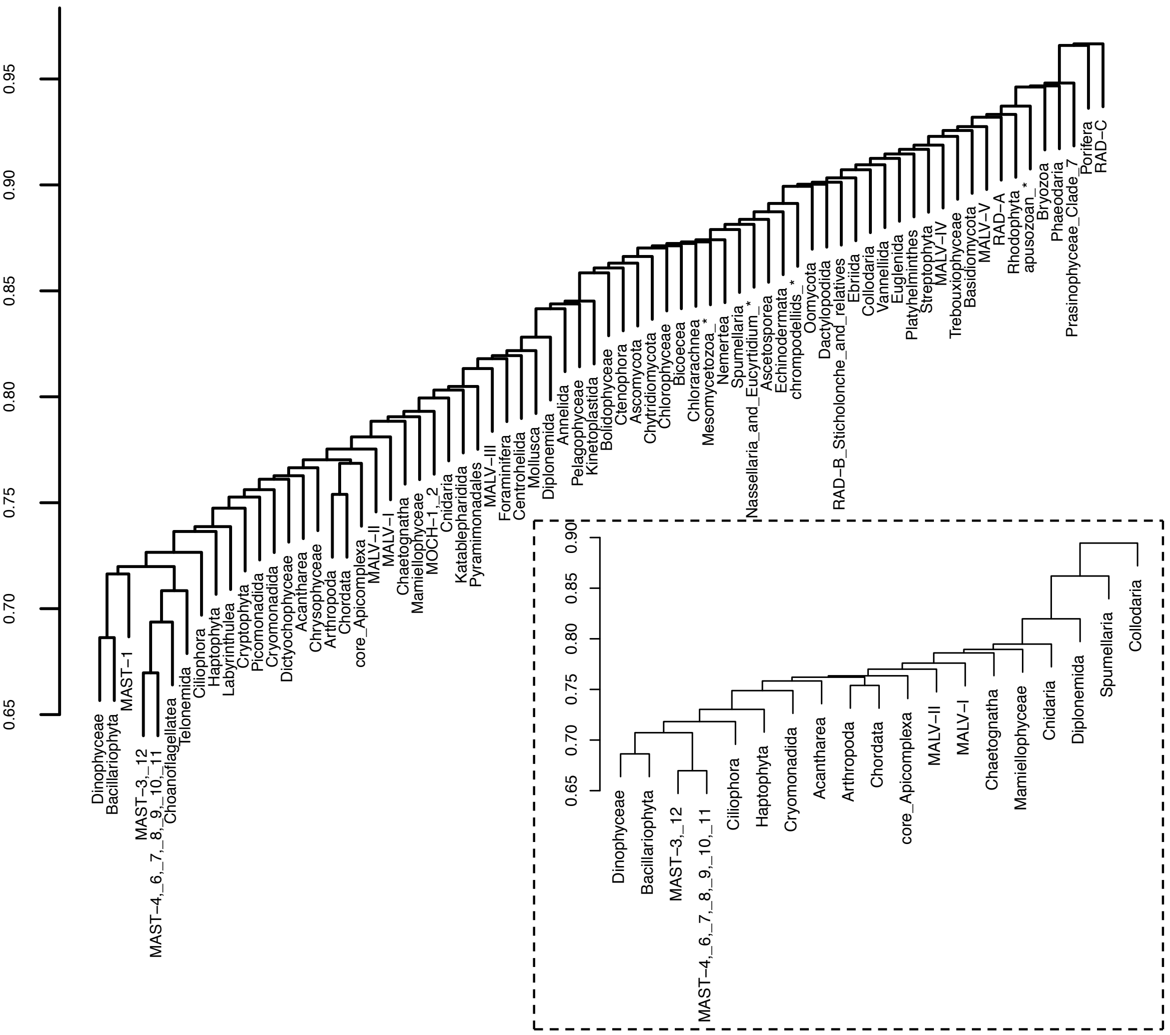


**Fig. S2. Dendrogram of the biogeographic dissimilarity between the 70 major groups of eukaryotic plankton.** Biogeographic dissimilarity was computed as the relative mutual information in the distribution of assemblages over stations. Inset: same plot, restricted to the 19 groups tallying more than 1,000 OTUs.

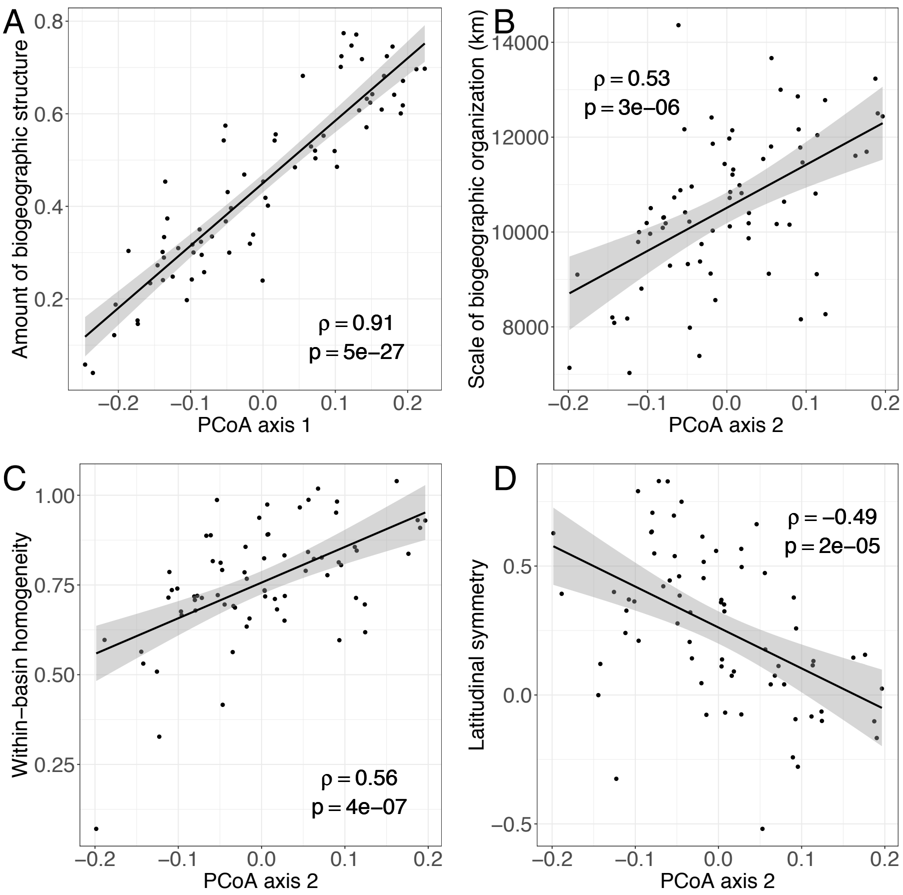


**Fig. S3. Interpretation of the first two axes of biogeographic variation based on surface spatial patterns.** (**A**) The amount of surface biogeographic structure displayed by each, as measured by the short-distance autocorrelation in the spatial distribution of assemblages at the surface, increases with group position on the first axis of biogeographic variation. (**B**) The scale of biogeographic organization at the surface, as measured by the characteristic distance at which spatial autocorrelation vanishes, increases with group position on the second axis. (**C**) The surface homogeneity of oceanic basins in terms of assemblages increases with group position on the second axis. (**D**) The symmetry of the surface assemblage distribution with respect to the Equator decreases with group position on the second axis.

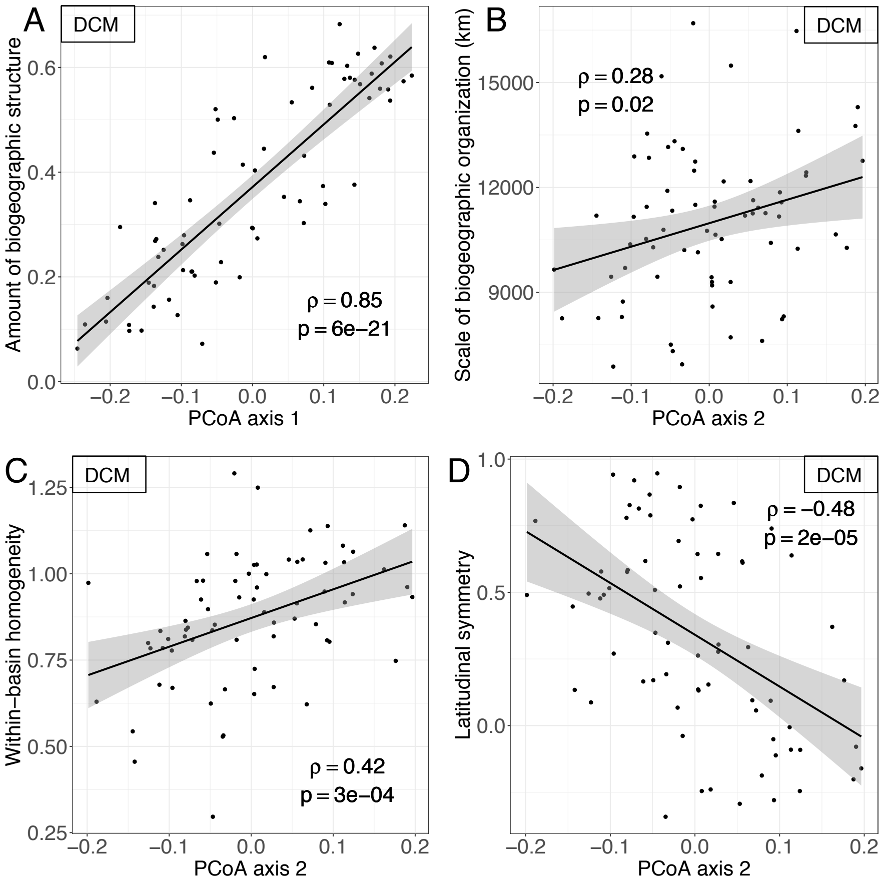


**Fig. S4. Interpretation of the first two axes of biogeographic variation based on DCM spatial patterns.** Same as Figure S3, but using DCM instead of surface results to compute (**A**) the amount of biogeographic structure, (**B**) the scale of biogeographic organization, (**C**) the homogeneity of oceanic basins in terms of assemblages, and (**D**) the symmetry of the assemblage distribution with respect to the Equator. The trends are the same as those observed at the surface, but weaker.

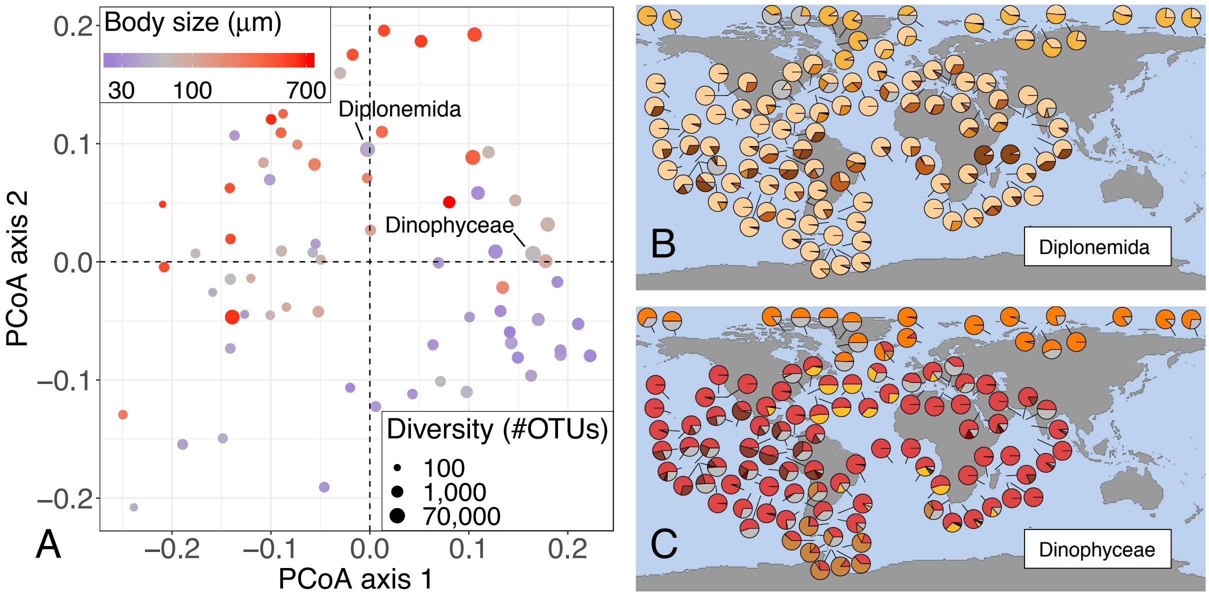


**Fig. S5. Surface biogeography of Diplonemida and Dynophyceae, the two most diverse groups of Eukaryotic plankton.**

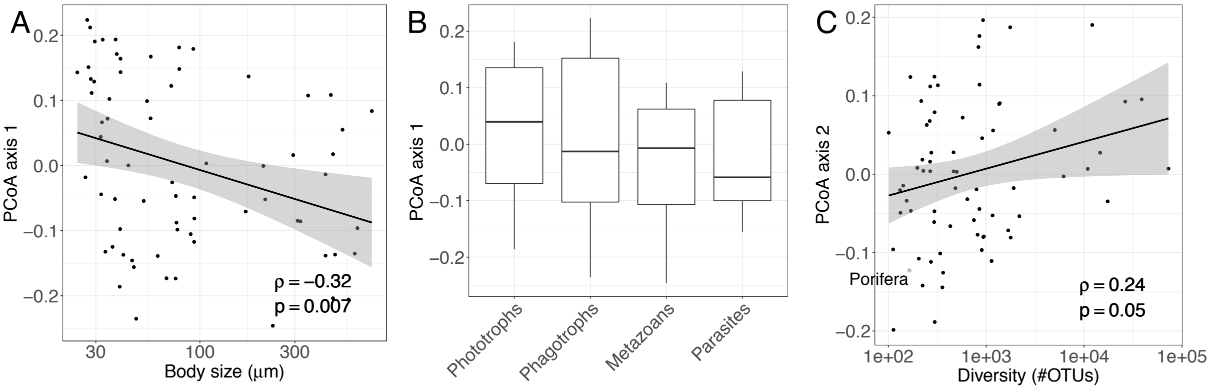


**Fig. S6. Relationship between biogeography and log diversity, log mean body size and ecology across major eukaryotic plankton groups – additional results with respect to Fig. 3.** (**A**) Group position along the first axis of biogeographic variation, indicative of the amount of biogeographic structure, versus log mean body size. (**B**) Differences in group position along the first axis between four broad ecological categories. Differences are not significant (ANOVA F-test $p = 0.62$), however phototrophs score significantly higher than metazoans once the dependence on diversity is taken into account (ANCOVA t-test: $p = 0.035$). (**C**) Group position along the second axis, indicative of the spatial scale and nature of the biogeographic structure, versus log diversity.

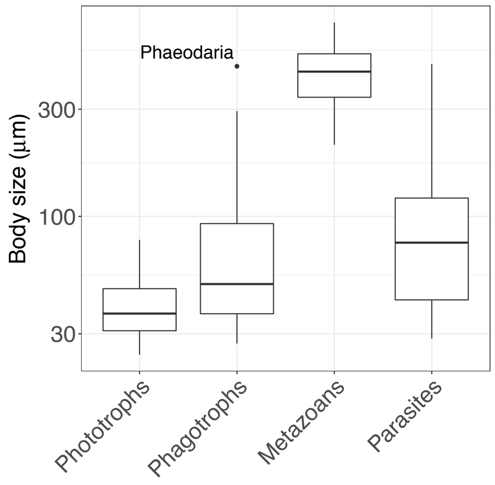


**Fig. S7. Body size variation across ecological categories.** Major eukaryotic plankton groups have significantly different mean log body sizes between the four ecological categories (ANOVA pairwise t-tests) except between phagotrophs and parasites.

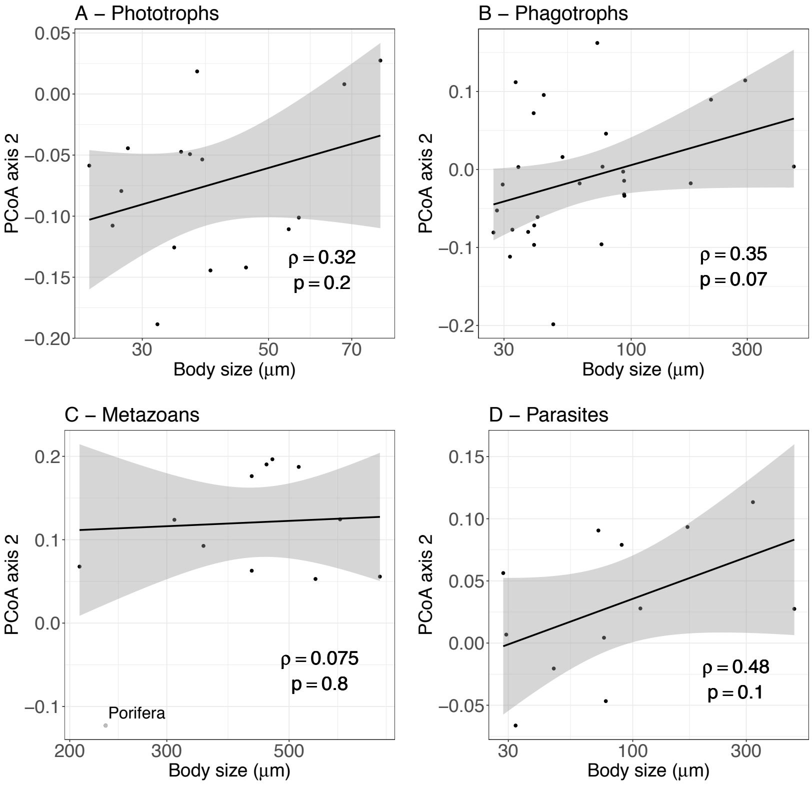


**Fig. S8. Group position on the second axis versus group mean body size within each ecological category.** The positive relationship between body size and position on the second axis still holds within each ecological category, even though the linear regressions are not statistically significant. An ANCOVA F-test with body size as covariate (excluding the outlier group Porifera) confirms this relationship ($p = 0.004$; $p = 10^{-4}$ if Metazoans and Parasites are grouped together).

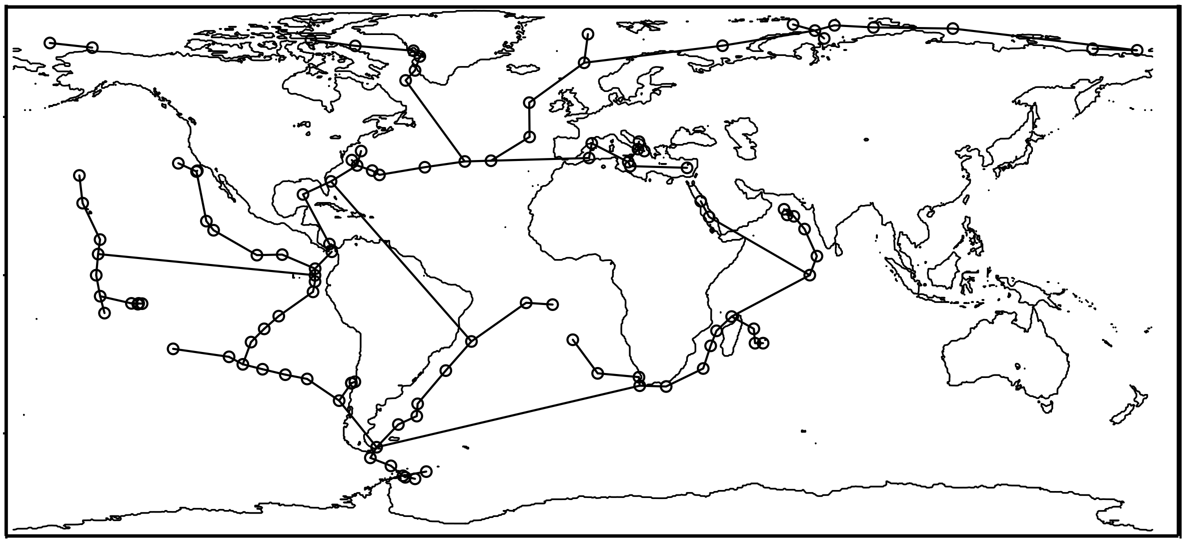

**Fig. S9. Minimal set of connections between all surface Tara stations.** Remaining connections between Tara stations after discarding all surface minimum travel times larger than 2.1 years. Discarding shorter minimum travel times leads to disconnecting some stations from the rest. We used this minimal set of connections between all surface Tara stations to construct connectivity maps by Principal Coordinate Analysis (Moran's Eigenvector Maps; cf. Mat. & Meth.).

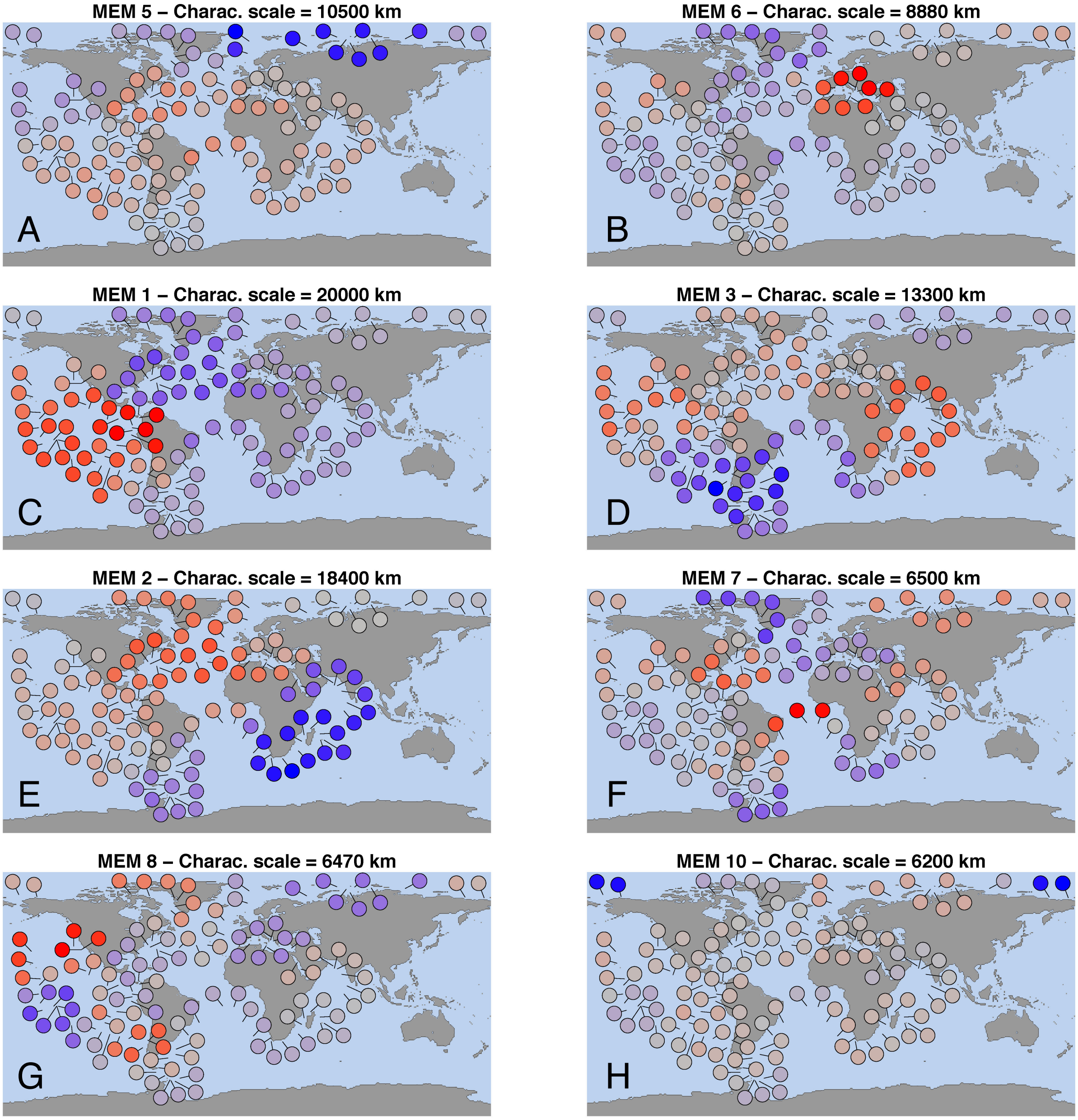


**Fig. S10. Connectivity maps most often selected across plankton groups.** (**A-H**) Moran Eigenvector Maps (MEMs), or connectivity maps, sorted by decreasing median amount of variance explained in the biogeography of plankton groups (see Fig. S13). Each MEM is identified by a number assigned by order of decreasing eigenvalue, and can be characterized by its characteristic scale of spatial autocorrelation (in km), computed using the same method as for the characteristic scale of biogeographic organization.

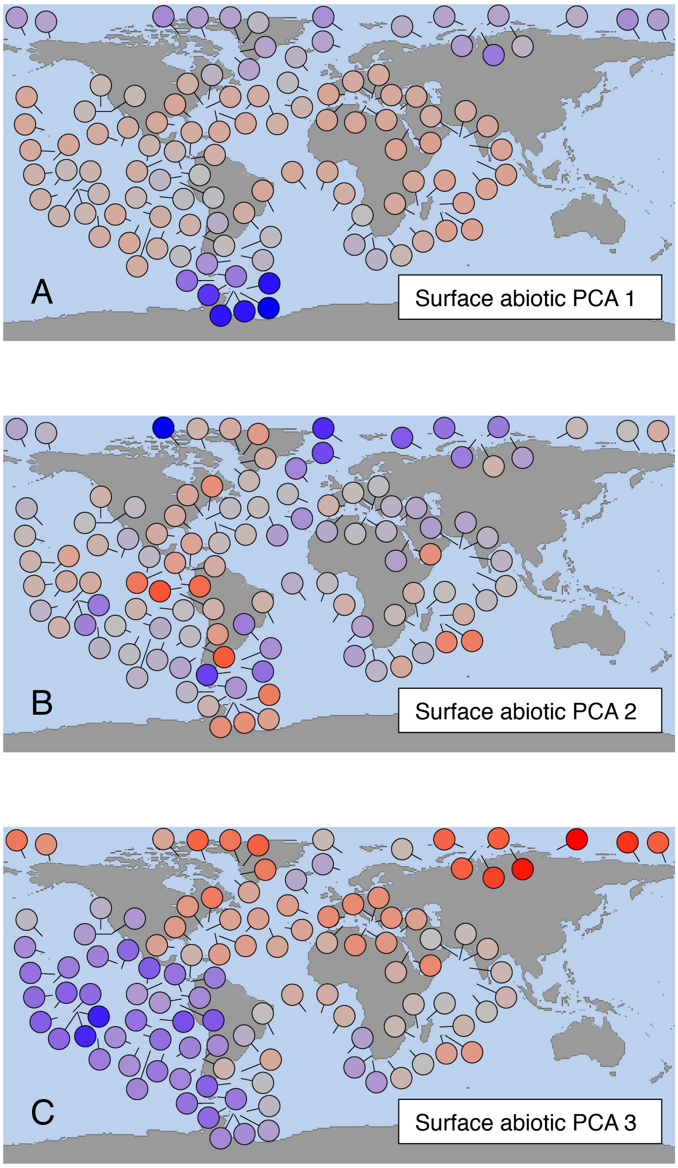


**Fig. S11. Surface abiotic environmental variables.** First three PCA axes for eight abiotic variables at the surface (temperature, nitrate, phosphate, silicate and iron concentrations, dissolved oxygen concentration, oxygen saturation and apparent oxygen utilization), used as abiotic explanatory variables for the biogeography of each taxonomic group. (**A**) The first PCA axis amounts to 44% of the variance and corresponds to variation in temperature as well as in nitrate, phosphate, silicate and dissolved oxygen concentrations. (**B**) The second PCA axis amounts to 26% of the variance and corresponds to variation in oxygen saturation and utilization. (**C**) The third PCA axis amounts 16% of the variance and corresponds to variation in iron concentration.

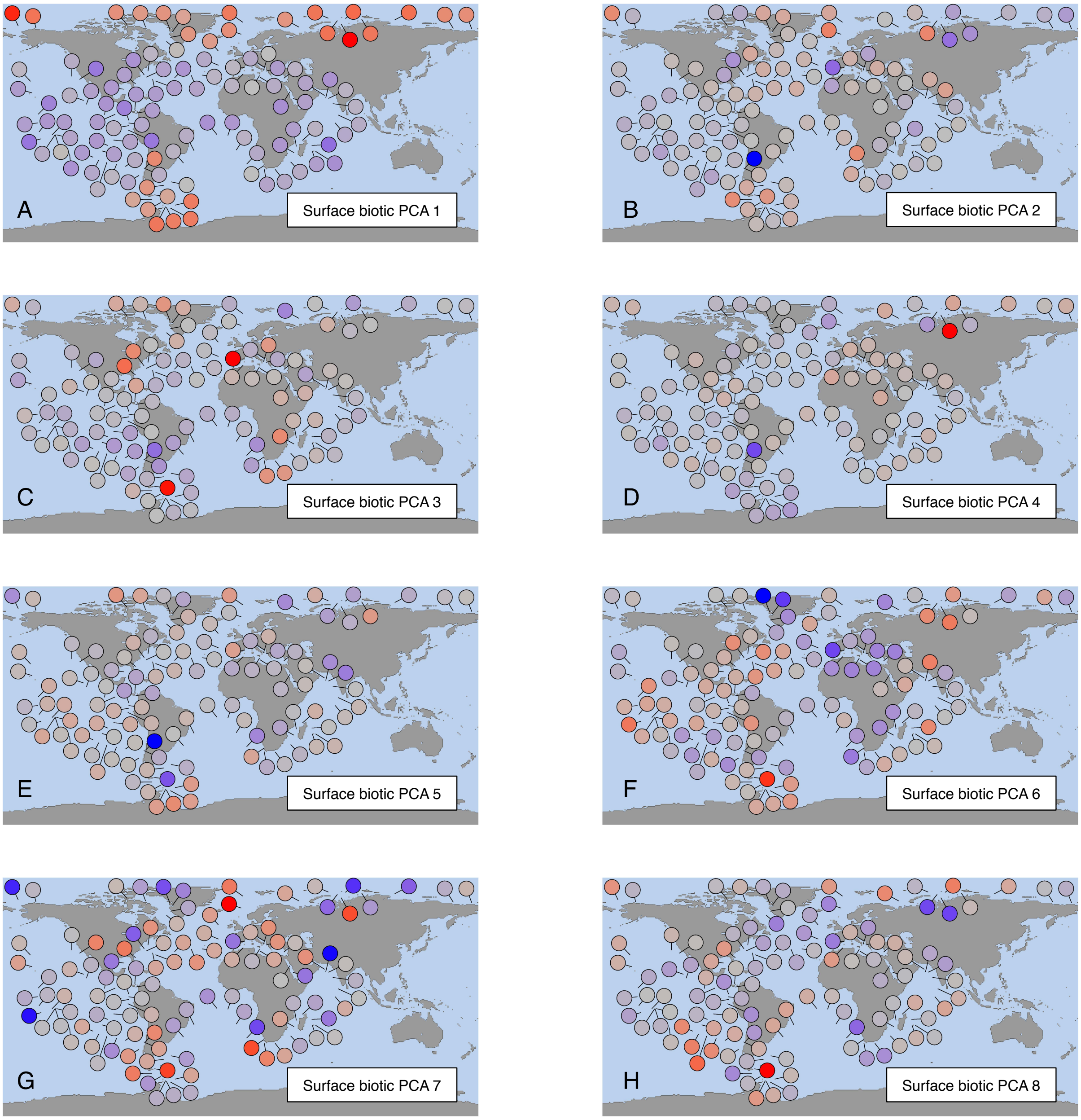


**Fig. S12. Surface biotic environmental variables.** First eight PCA axes for the surface relative abundance of the 70 major eukaryotic groups across stations, estimated based on their relative read counts. We used the first twenty-eight axes as biotic explanatory variables for the biogeography of each group, after removing the focal group from the PCA. The first eight PCA axes account for (**A**) 10.5%, (**B**) 6.1%, (**C**) 5.3%, (**D**) 5.0%, (**E**) 4.7%, (**F**) 4.3%, (**G**) 3.5% and (**H**) 3.4% of the variance, respectively.

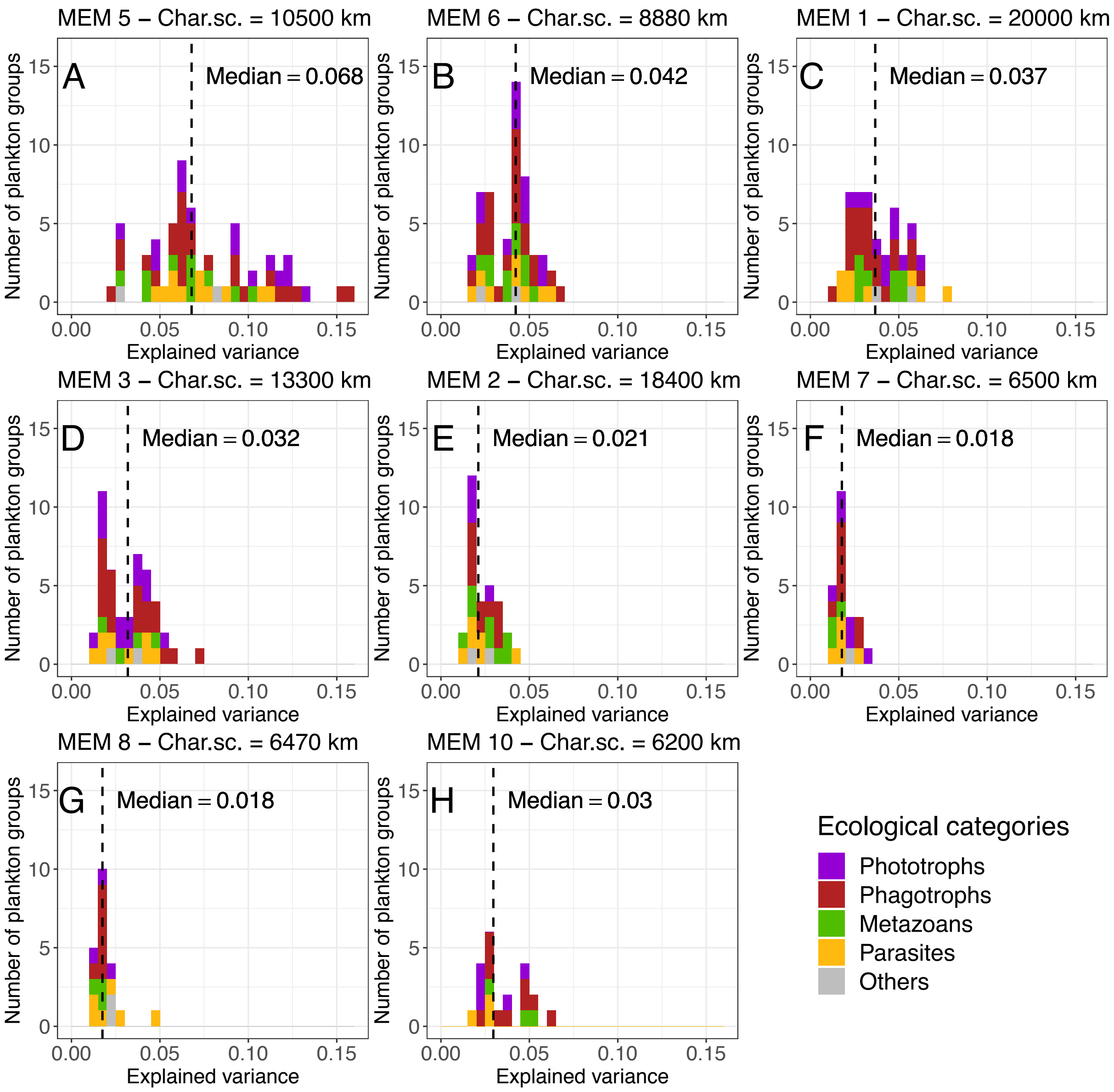


**Fig. S13. Variance explained by the eight connectivity maps that were most often selected across plankton groups.** (**A-H**) For each Moran Eigenvector Map (MEM), histogram of the variance explained in the biogeography of plankton groups, across groups for which the MEM has been retained by the variable selection procedure. MEMs are ordered as in Figure S10 by decreasing median amount of variance explained in the biogeography of plankton groups, shown by a dashed line. The most selected eigenvectors correspond to connectivity patterns ranging from the basin scale (about 6,000 km) to the global scale (about 20,000 km); see Fig. S10 for the corresponding spatial representations. The ecological categories of the selected groups are indicated by colors. There is not significant difference across ecological categories except for MEM 2 (which explains more variance for metazoans than phototrophs) and MEM 5 (the reverse).

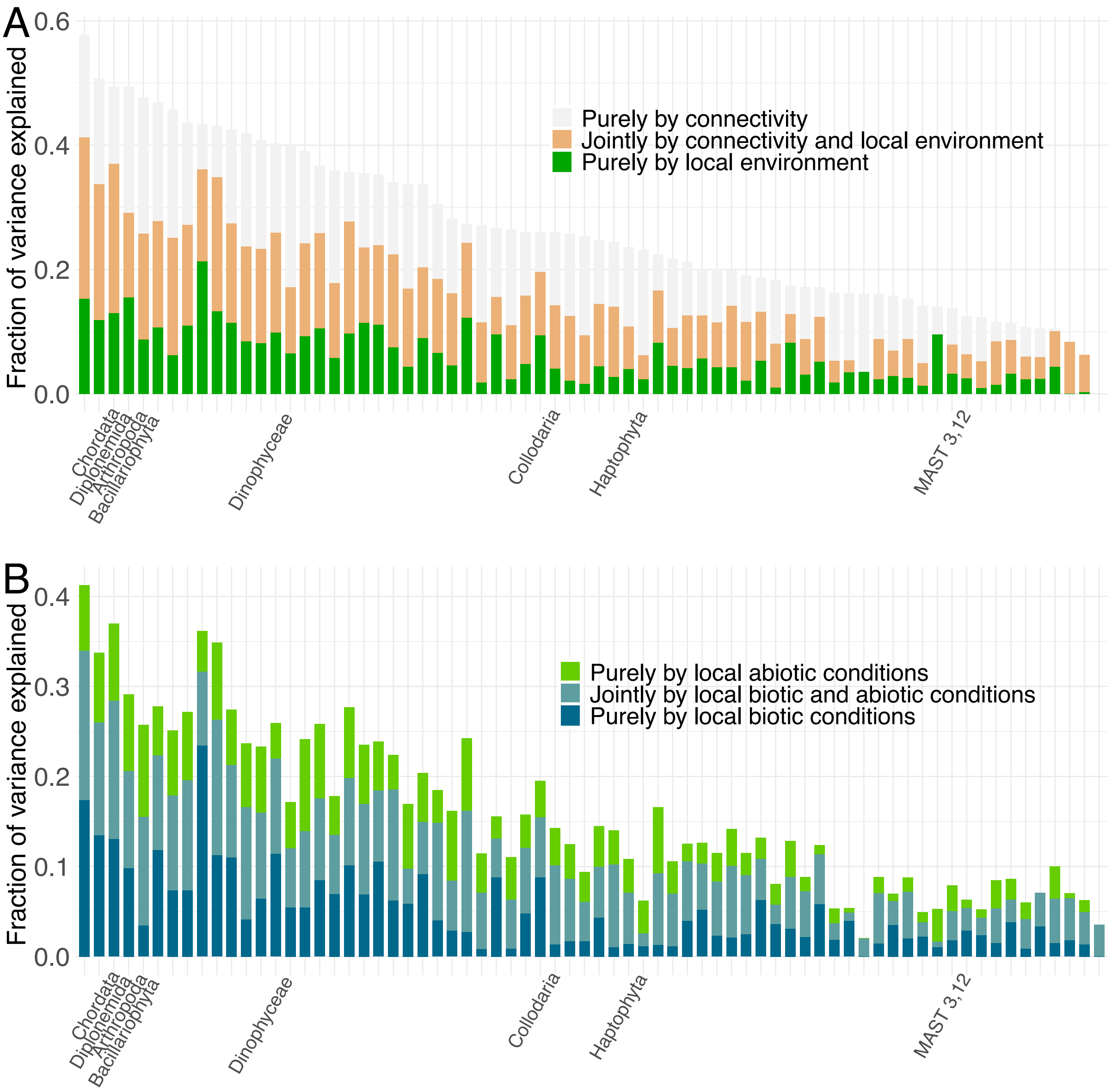


**Fig. S14. Variance in the surface biogeography of major eukaryotic plankton groups that can be explained by connectivity maps, the local abiotic environment or the local biotic environment.** (**A**) Fractions of variance that can be explained purely by connectivity maps (upper fraction), jointly by connectivity maps and the (biotic and abiotic) local environment (middle fraction) and purely by the local environment (lower fraction), for all major taxonomic groups ordered by decreasing total explained variance (excluding Porifera, for which no explanatory variable was selected). (**B**) Within the variance that can be explained by the (biotic and abiotic) local environment, fractions of variance that can be explained purely by abiotic conditions (upper fraction), jointly by abiotic and biotic conditions (middle fraction) and purely by biotic conditions (lower fraction) for the groups ordered as in panel (A).

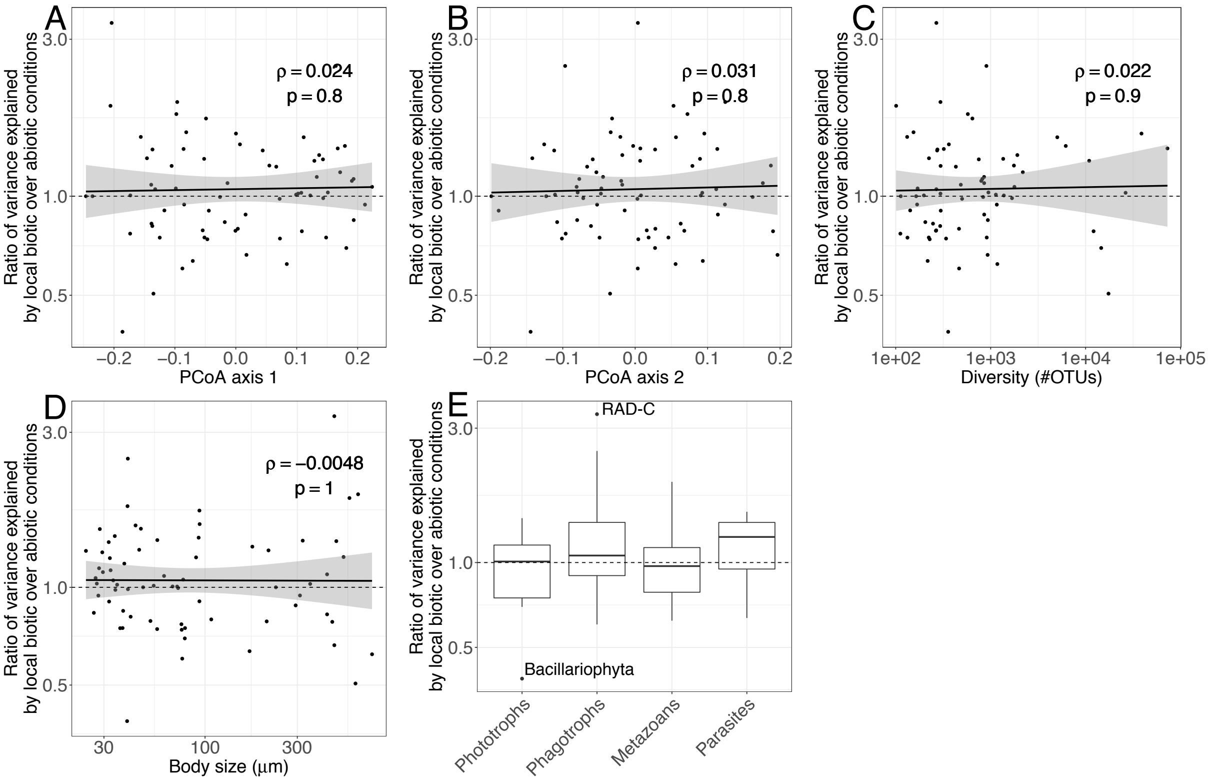


**Fig. S15. Ratio of the variance explained by biotic and abiotic environmental conditions in the surface biogeography of major taxonomic groups.** Log ratio of the variance explained by biotic conditions over the variance explained by abiotic conditions across major taxonomic groups, versus (**A**) group position on the first axis of variation, (**B**) group position on the second axis of variation, (**C**) group diversity, (**D**) group mean body size and (**E**) between broad ecological categories. The ratio is distributed around 1, reflecting an approximately equal influence of biotic and abiotic conditions overall. It does not significantly vary with group position on the first two axes of biogeographic variation, diversity, body size or ecology.

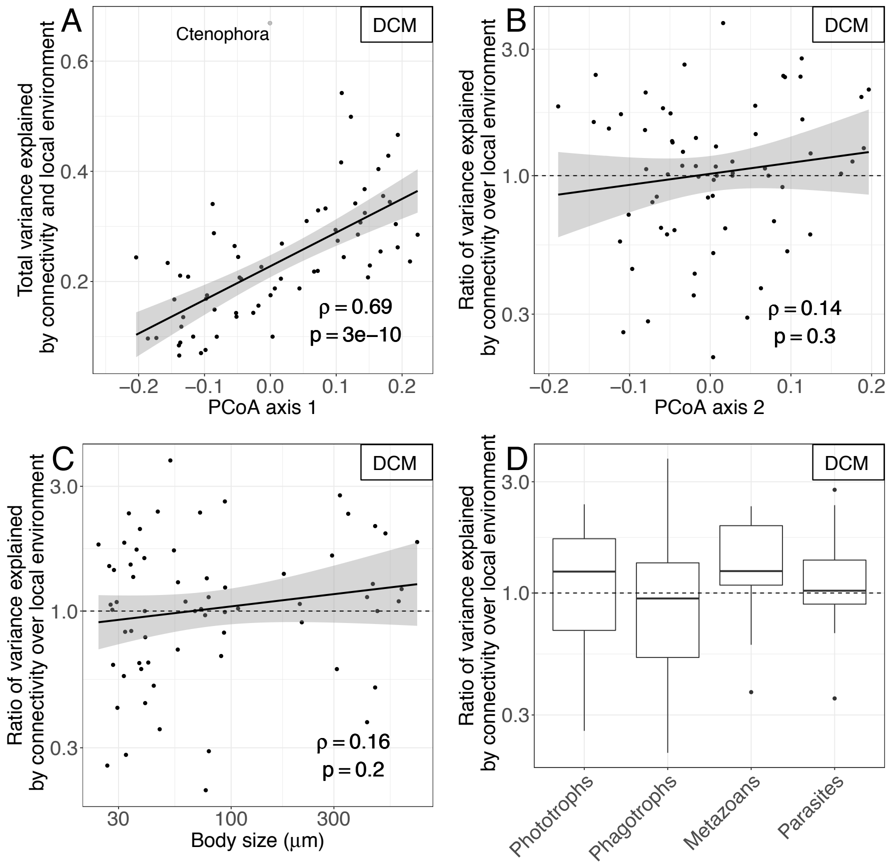


**Fig. S16. Drivers of DCM biogeography across major eukaryotic plankton groups.** Same as Figure 4, but computed at the DCM. The trends are qualitatively the same, but they are much weaker and only (**A**) the positive relationship between the total explained variance and the group position on the first axis of biogeographic variation is significant. (**B-D**) The ratio of the variance explained purely by connectivity maps and by the (biotic and abiotic) local environment is closer to one than at the surface. This may be partly due do the fact that the estimated travel times along currents at the DCM are not as good an approximation to connectivity as at the surface, since they were simulated at 75 m depth whereas the actual DCM samples were collected over a range of depths. The grey dot denotes Ctenophora, an outlier group excluded from statistical tests.

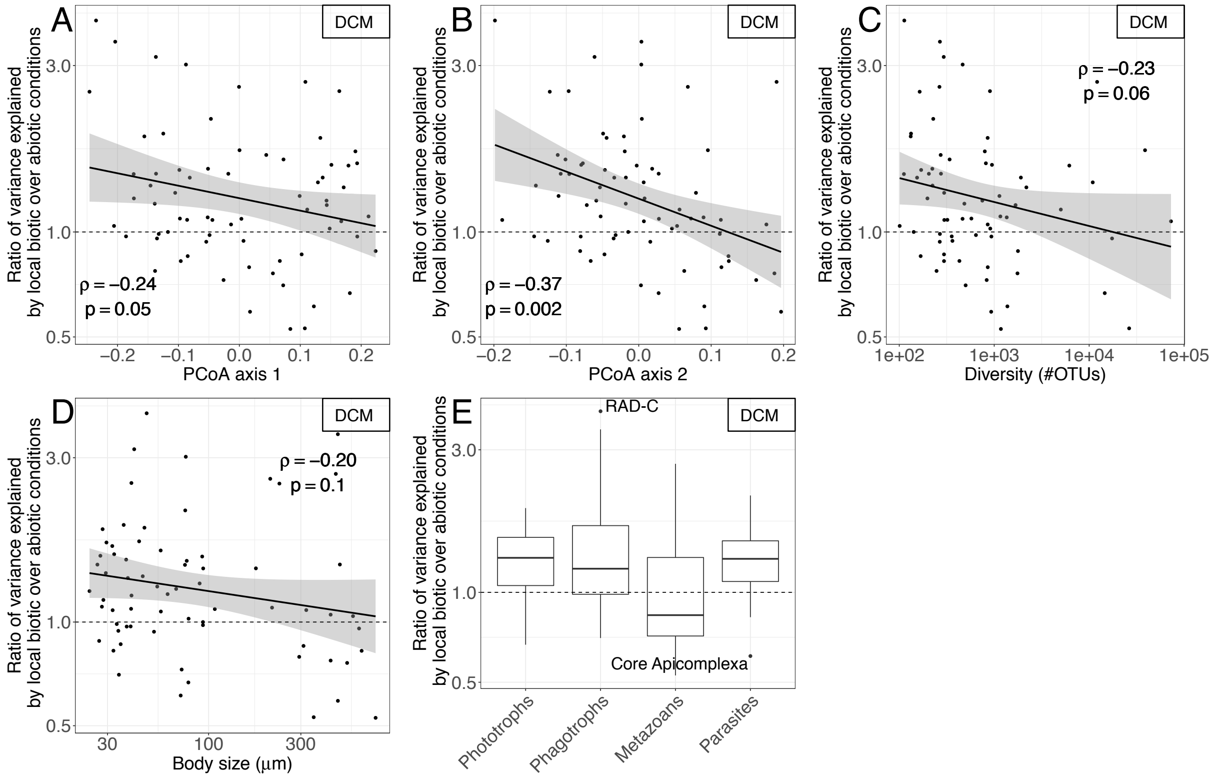


**Fig. S17. Ratio of the variance explained by biotic and abiotic environmental conditions in the DCM biogeography of major taxonomic groups.** Same as Figure S15, but computed at the DCM. As at the surface, the ratio of the variance explained purely by biotic conditions and purely by abiotic conditions does not significantly vary with group position on the first two axes of biogeographic variation, diversity, body size or ecology.

| *Clade* | *Diversity (#OTUs)* | *Body size (μm)* | *Dominant ecology* | *PCoA 1* | *PCoA 2* | *LDA stability across runs* |
|---|---|---|---|---|---|---|
| Dinophyceae | 72769 | 57 | NA | 0.16 | 0.01 | 0.90 |
| Diplonemida | 38769 | 44 | Phagotrophs | 0.00 | 0.10 | 0.80 |
| Arthropoda | 26366 | 350 | Metazoans | 0.10 | 0.09 | 0.85 |
| Collodaria | 17417 | 600 | NA | -0.14 | -0.05 | 0.87 |
| Bacillariophyta | 14592 | 79 | Phototrophs | 0.18 | 0.03 | 0.90 |
| Chordata | 12129 | 455 | Metazoans | 0.11 | 0.19 | 0.78 |
| MALV-II | 10909 | 29 | Parasites | 0.13 | 0.01 | 0.90 |
| Ciliophora | 6179 | 93 | Phagotrophs | 0.18 | 0.00 | 0.86 |
| MALV-I | 5031 | 29 | Parasites | 0.11 | 0.06 | 0.83 |
| Haptophyta | 2182 | 38 | Phototrophs | 0.17 | -0.05 | 0.86 |
| Acantharea | 1910 | 176 | Phagotrophs | 0.13 | -0.02 | 0.80 |
| MAST-3, 12 | 1779 | 27 | Phagotrophs | 0.22 | -0.08 | 0.87 |
| Cnidaria | 1758 | 520 | Metazoans | 0.05 | 0.19 | 0.75 |
| Cryomonadida | 1678 | 40 | Phagotrophs | 0.14 | -0.07 | 0.85 |
| Apicomplexa | 1377 | 72 | Parasites | 0.12 | 0.09 | 0.79 |
| Spumellaria | 1355 | 213 | Phagotrophs | -0.06 | 0.08 | 0.79 |
| Chaetognatha | 1176 | 731 | Metazoans | 0.08 | 0.05 | 0.82 |
| MAST-4, 6, 7, 8, 9, 10, 11 | 1161 | 28 | Phagotrophs | 0.21 | -0.05 | 0.80 |
| Mamiellophyceae | 1135 | 54 | Phototrophs | 0.10 | -0.11 | 0.77 |
| Dictyochophyceae | 947 | 28 | Phototrophs | 0.15 | -0.08 | 0.76 |
| Choanoflagellatea | 934 | 38 | Phagotrophs | 0.19 | -0.08 | 0.80 |
| Mollusca | 929 | 466 | Metazoans | 0.01 | 0.20 | 0.77 |
| Picomonadida | 909 | 79 | Phagotrophs | 0.15 | 0.05 | 0.76 |
| Labyrinthulea | 902 | 40 | Phagotrophs | 0.16 | -0.10 | 0.77 |
| Foraminifera | 855 | 294 | Phagotrophs | 0.01 | 0.11 | 0.78 |
| Annelida | 854 | 428 | Metazoans | -0.02 | 0.18 | 0.75 |
| Chrysophyceae | 853 | 28 | Phototrophs | 0.13 | -0.04 | 0.74 |
| Ascomycota | 833 | 73 | Phagotrophs | -0.03 | 0.16 | 0.58 |
| Telonemida | 817 | 32 | Phagotrophs | 0.19 | -0.07 | 0.76 |
| MAST-1 | 796 | 30 | Phagotrophs | 0.19 | -0.02 | 0.73 |
| Cryptophyta | 750 | 24 | Phototrophs | 0.14 | -0.06 | 0.76 |
| Nassellaria & Eucyrtidium | 642 | 94 | Phagotrophs | -0.05 | -0.04 | 0.70 |
| RAD-B (Sticholonche & relatives) | 575 | 40 | Phagotrophs | -0.10 | 0.07 | 0.65 |
| Katablepharidida | 500 | 34 | Phagotrophs | 0.07 | 0.00 | 0.64 |
| Basidiomycota | 485 | 61 | Phagotrophs | -0.14 | -0.01 | 0.47 |
| Ebriida | 465 | 76 | Phagotrophs | -0.09 | 0.01 | 0.64 |
| Kinetoplastida | 464 | 108 | Parasites | 0.00 | 0.03 | 0.65 |
| MALV-III | 430 | 32 | Parasites | 0.06 | -0.07 | 0.64 |
| Pelagophyceae | 363 | 34 | Phototrophs | 0.01 | -0.12 | 0.71 |
| Prasinophyceae Clade 7 | 357 | 40 | Phototrophs | -0.19 | -0.15 | 0.61 |
| Pyramimonadales | 339 | 57 | Phototrophs | 0.07 | -0.10 | 0.75 |
| Ascetosporea | 321 | 320 | Parasites | -0.09 | 0.11 | 0.55 |

| | | | | | | |
|---|---|---|---|---|---|---|
| Chlorophyceae | 298 | 32 | Phototrophs | -0.05 | -0.19 | 0.68 |
| Oomycota | 297 | 90 | Parasites | -0.11 | 0.08 | 0.54 |
| Echinodermata | 296 | 619 | Metazoans | -0.10 | 0.12 | 0.66 |
| MOCH-1, 2 | 296 | 35 | Phototrophs | 0.10 | -0.05 | 0.70 |
| RAD-A | 294 | 41 | Phagotrophs | -0.14 | -0.07 | 0.52 |
| MALV-IV | 275 | 477 | Parasites | -0.14 | 0.02 | 0.49 |
| Centrohelida | 273 | 32 | Phagotrophs | 0.04 | -0.11 | 0.69 |
| Phaeodaria | 268 | 466 | Phagotrophs | -0.21 | 0.00 | 0.56 |
| Vannellida | 268 | 34 | Phagotrophs | -0.14 | 0.11 | 0.60 |
| Bicoecea | 267 | 52 | Phagotrophs | -0.06 | 0.01 | 0.68 |
| Ctenophora | 265 | 208 | Metazoans | 0.00 | 0.07 | 0.60 |
| Platyhelminthes | 247 | 428 | Metazoans | -0.14 | 0.06 | 0.50 |
| Mesomycetozoa | 228 | 76 | Parasites | -0.05 | 0.00 | 0.54 |
| Trebouxiophyceae | 224 | 46 | Phototrophs | -0.15 | -0.15 | 0.54 |
| Chlorarachnea | 223 | 37 | Phototrophs | -0.05 | 0.02 | 0.62 |
| Chytridiomycota | 217 | 170 | Parasites | -0.07 | 0.10 | 0.52 |
| Bolidophyceae | 205 | 27 | Phototrophs | -0.02 | -0.11 | 0.60 |
| Rhodophyta | 198 | 68 | Phototrophs | -0.18 | 0.01 | 0.44 |
| Dactylopodida | 170 | 77 | Parasites | -0.10 | -0.05 | 0.58 |
| Nemertea | 168 | 310 | Metazoans | -0.09 | 0.13 | 0.54 |
| Porifera | 164 | 232 | Metazoans | -0.25 | -0.13 | 0.32 |
| Chrompodellids | 154 | 94 | Phagotrophs | -0.08 | -0.04 | 0.46 |
| Euglenida | 142 | 94 | Phagotrophs | -0.12 | -0.01 | 0.52 |
| Streptophyta | 133 | 36 | Phototrophs | -0.13 | -0.04 | 0.45 |
| MALV-V | 132 | 47 | Parasites | -0.16 | -0.03 | 0.42 |
| RAD-C | 113 | 48 | Phagotrophs | -0.24 | -0.21 | 0.31 |
| Bryozoa | 101 | 558 | Metazoans | -0.21 | 0.05 | 0.37 |

**Table S1. Major eukaryotic clades sorted by decreasing diversity.** Columns indicate the number of OTUs assigned to the clade, the mean body size for the clade estimated from sampling size fractions, the ecological category assigned to the clade based on its known dominant ecology, the projection of the clade onto the first two axes of biogeographic variation, and the robustness of the Latent Dirichlet Allocation performed on the clade, as measured by the stability of the biogeography obtained across random initial conditions for the inference algorithm.
